# Seqrutinator: Non-Functional Homologue Sequence Scrutiny for the Generation of large Datatsets for Protein Superfamily Analysis

**DOI:** 10.1101/2022.03.22.485366

**Authors:** Agustín Amalfitano, Nicolás Stocchi, Hugo Marcelo Atencio, Fernando Villarreal, Arjen ten Have

## Abstract

**Background:** In recent years protein bioinformatics has resulted in many good algorithms for multiple sequence alignment (MSA) and phylogeny. Little attention has been paid to sequence selection whereas notably recently published complete proteomes often have many sequences that are partial or derive from pseudogenes. Not only do these sequences add noise to the MSA, phylogeny and other downstream computational analyses, they also instigate many errors in the processing of the MSAs and downstream analyses, including the phylogeny.

**Objective:** This work aims to provide and test an objective, automated but flexible pipeline for the scrutiny of sequence sets from large, complex, eukaryotic protein superfamilies. The pipeline should classify sequences with high precision and recall as either functional or non-functional. The pipeline should classify no or only a few SwissProt sequences as non-functional (high precision) and sequences from other related superfamilies as non-functional (high recall) and result in a demonstrably much improved MSA (high performance).

**Results:** Seqrutinator is a pipeline that consists of five modules written in Python3 that identify and remove sequences that are likely Non-Functional Homologues (NFH). Here we tested the pipeline using three complex plant superfamilies (BAHD, CYP and UGT) that act in specialized metabolism, using the complete proteomes of 16 plant species as input and SwissProt as a control. Only 1.94% of SwissProt sequences with wetlab evidence were identified as NFH and all sequences from other related superfamilies were removed. Most NFH sequences are partial but, interestingly, their removal results in highly improved MSAs. a few but significant sequences that instigate large gaps were found. The five modules show similar behaviour when applied to the 16 sequence sets of the three analysed superfamilies. Pipelines with different module orders result in similar classifications and, moreover, show that different modules often detect the same sequences.

**Conclusion and perspective:** Seqrutinator forms a consistent pipeline for sequence scrutiny that does result in sequence sets that generate high fidelity MSAs. Recovery analyses show the method has high precision and recall.

## Introduction

Protein superfamilies, here defined as protein families with subfamilies that have different functional characteristics, are the subject of many computational studies (e.g. [1–8]) and form the target of many computational platforms (e.g. [9–12]). Structure-function analysis aims not only to identify which residues and/or subsequences are involved in functional diversification, it also tries to explain and predict the functional differences and can identify hitherto nondescript subfamilies (e.g. [2, 3]). A large set of methods (e.g. [13–15]) is available and novel methods are published regularly (for review see [16]) in a research field often referred to as phylogenomics.

The basis for many protein bioinformatics tools is formed by phylogenies and their underlying Multiple Sequence Alignments (MSAs). Hence, a large set of algorithms and programs has been developed for both. Reliable methods for superfamily phylogeny use maximum likelihood (PHYML [17], RAXML [18]; FASTTREE [19]) or Bayesian inference (MrBayes [20]). The construction of MSAs has improved significantly in recent years [21–25] but, likely since more complex protein families are now being analysed, is also still one of the major areas of research in bioinformatics (e.g. [26]).

Only recently, attention has been paid to automated sequence scrutiny, the selection of sequences that are to be included in the analysis [27–31]. This is a laborious task where a high complexity of a protein family hampers the identification of Non-Functional Homologues (NFHs). Information in an MSA provided by sequences that do not correspond to a Functional Homologue (FH) is often considered as mere noise that in all likelihood does not have a significant effect on the outcome of analyses. MSAs are therefore often trimmed in order to remove columns with low reliability [32, 33]. We argue that, besides that this is an actual loss of information, incorrect sequences of NHFs can also prevent or hinder the correct processing of the MSA, thereby generating erroneous signal.

There are two major sources of NFH sequences: Pseudogenes and incorrect sequences. Pseudogenes show similarity levels from low (i.e. non detectable) to high (i.e. indistinguishable from FHs). Since they are no longer under functional constraint they can accumulate not only point mutations but also obtain inserts and or deletions of subsequences, especially when the original gene contained introns. Incorrect sequences can, in their turn, result from both sequencing errors and incorrect gene models. Notably recently published complete proteomes often contain many incorrect gene models.

Datasets that consist of sequences from many complete proteomes are often prohibitively long due to an accumulation of various errors. E.g. sequencespecific inserts provide information that can derail proper MSA. Sequence scrutiny is therefore required but this demands a huge effort when large datasets are studied. Existing methods are either not fully objective [27], directed at improving existing MSAs by removing subsequences [29], or only remove outliers [28, 30, 31]. None of these methods is fully automated and directed at removing NFH sequences from large sequence sets in order to obtain clean datasets. Most of the existing algorithms are only tested on simulated datasets and none of them directs the problem of large gap regions. This is likely caused by the fact that defining inclusion thresholds is troublesome and will almost by definition result in both false positives and false negatives. In addition, no real benchmark datasets of FHs exist and any attempt to construct a benchmark set will result in a set that is too restrictive.

We developed a method for objective sequence scrutiny named Seqrutinator, directed at the removal of sequences from NFHs. As such, with scrutiny, we refer to the identification and removal of NFH sequences. The method was developed and tested by performing the sequence mining of three of the most complex single domain superfamilies in plants, to wit cytochrome P450 (CYP), UDP-Glycosyl Transferase (UGT), and BAHD acyltransferases (BAHD). Interestingly, many enzymes of these superfamilies function in specialized metabolism by which these families show a high evolutionary rate. This explains in part the high diversity and complexity of these superfamilies and makes sequence scrutiny of these families hard. The case studies were used to improve the procedure with the final objective of constructing an automated pipeline. Here we present the pipeline, separate scripts and methods. We show numerical data generated by the three case studies directed at the validation of the method. We show that the MSAs of scrutinized datasets are significantly more reliable and that the method is flexible and robust. Most importantly, Seqrutinator appears to remove mostly real NFHs, it has only a few false positives (FHs classified as NFH) whereas no false negatives (NFHs classified as FH) were detected. Furthermore, we present a detailed recovery analysis that, besides the high performance of Seqrutinator, shows the relative ease of sequence analysis once a high-quality MSA has been obtained.

### Design of the pipeline and its modules

#### Objective

We define the objective of this work as to provide and test an objective, automated but flexible pipeline for the scrutiny of sequence sets for large, complex eukaryotic protein superfamilies. Conceptually, the method classifies sequences as either functional or non-functional. Input sequence sets represent all identified probable homologues of a protein superfamily, obtained by a sensitive data-mining methodology. Output sets consist of the aligned sequence set of FHs and the NFH sequence set. The FH set should be aligned without much uncertainty. The NFH set should be subjected to a recovery analysis in order to prevent inadvertent false positives (FHs classified as NFH). The method should not be seen as a method to predict whether a sequence is functional or not. It merely removes many sequences that likely correspond to NFHs with the objective to obtain a high-quality MSA representing a protein superfamily.

#### Definition of Non-Functional Homologue

To scrutinize protein sequence datasets for NFHs, we need to define NFH in terms of sequence characters. We describe two major classes of NFH sequences that are further subdivided. First, NFH sequences can result from either incorrect gene modelling or sequencing errors. Second, NFH sequences can correspond to pseudogenes, where we define a pseudogene as a gene that no longer encodes its supposed or original function. As such, the scrutiny we develop is (super)family-dependent.

Incorrect gene modelling and sequencing errors can result in several issues. First, some sequences will lack either N- or C-terminal subsequences as a result of a missed start and/or stop codon. Other sequences are incomplete as a result of incomplete sequencing or incorrect assembly. Second, missed start and stop codons can also lead to additional subsequences at either the N- or C-terminus. Third, not all introns are identified from eukaryotic sequences. Coding subsequences that are incorrectly identified as intron form the fourth problem. Both these intron issues can lead to intron-sized gaps or a switch in the reading frame and the untimely stop of the coding sequence.

Pseudogenes are no longer under constraint and will as such accumulate mutations. This can result in two scenarios. It will result in an increased sequence and evolutionary distance, as can often be seen in phylogenetic trees. It may also result in the loss or gain of start and stop codons as well as splice donor and acceptor sites. As such, many pseudogenes will have issues similar to those found in NFH sequences that result from incorrect gene modelling or sequencing errors. The last issue is that depending on the sensitivity of the initial data mining and the possible existence of homologous superfamilies, certain initially identified sequences may be functional but do not belong to the superfamily of interest. For simplicity, we consider these also as NFH.

Based on the above problem description we hypothesize that:

1. Relatively short sequences are unlikely functional.
2. The presence of large continuous regions of gap-rich columns in MSAs can be instigated by one or more non-functional sequences with intron-derived subsequences incorrectly called as exons during gene modelling.
3. The presence of large continuous gaps in aligned sequences at otherwise amino acid-rich columns can occur in non-functional sequences due to the absence of exon-derived subsequences incorrectly called as intron during gene modelling.
4. Low similarity of a sequence to a HMMER profile suggests the sequence does not belong to the protein family presented by the HMMER profile, being either not functional or having a different function.

Modules and algorithms will be designed to find NFH sequences based on these four hypotheses. Note that the resulting method, named Seqrutinator, is based on the concept of homology which makes Seqrutinator principally not a valid method for protein families with different domain architectures.

#### The Modules and the Default Pipeline

The fully automated Seqrutinator pipeline is implemented in a larger procedure that starts with a user-guided sequence collection and ends with user-guided recovery analyses, as shown in Figure 1 and described in more detail in Supplemental Document 1. Sequence collection from multiple sequence sets is fully automated using a simple script named Multiple FASta Aligner (MUFASA, see also Supplemental Document 1). This requires a HMMER profile as well as initial sequence sets (e.g. complete proteomes) as input. All sequences from a complete proteome with hits against the HMMER profile are collected in a single fasta file and aligned automatically. A reference sequence selected by the user is aligned to each MSA using MAFFT add in order to remove non-homologous N- and C-terminal subsequences which may negatively interfere with the automated procedure.

**Figure 1:**
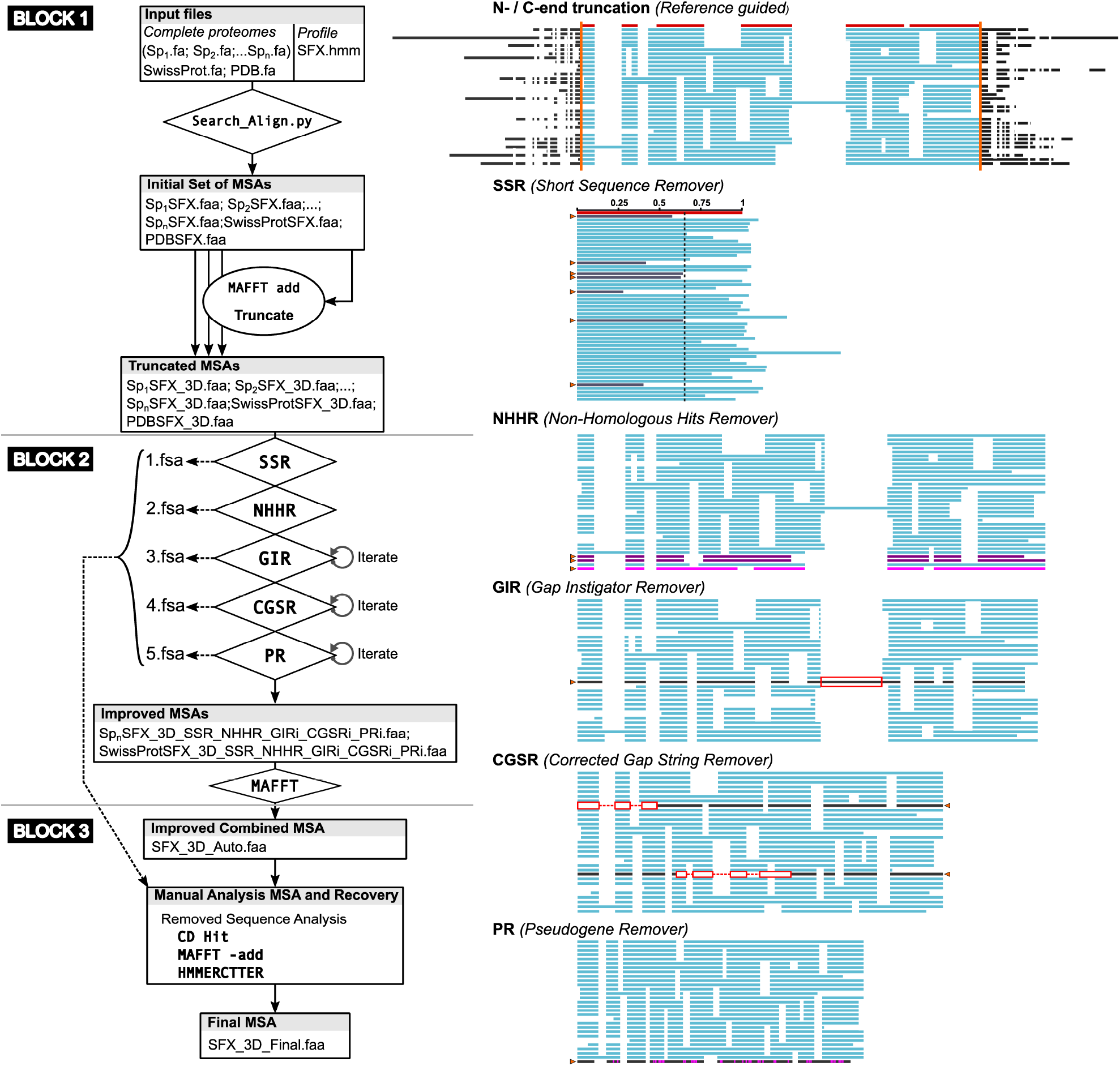
Schematic of the Default Seqrutinator Pipeline. The complete workflow for protein superfamily sequence mining consists of three blocks (left). The first block concerns the preparation of the input for the automated Seqrutinator pipeline in the second block. Each of Seqrutinators modules, here shown in its default order, and its iterations end with re-alignment of the remaining sequences. The third block concerns analyses for the recovery of inadvertently removed FH sequences. 1.fsa, 2.fsa, 3.fsa, 4.fsa and 5.fsa are archives in which the removed sequences of the subsequent pipeline modules are temporarily stored for recovery analyses. The schematic MSAs on the right show how the truncation of block 1, and how the modules of the automated pipeline in block 2, function.

Seqrutinator is a flexible pipeline made of five different modules. The user can select the modules and their order and change settings that will affect the stringency of the automated scrutiny. Below we describe the default procedure (see Figure 1) and its reasoning. Table 1 shows which defective sequences may be detected by which modules. The pipeline always starts with a single script that removes all sequences with non-IUPAC code. All other details are in the detailed description of the pipeline and modules in Supplemental Document 1.

**Table 1:** Seqrutinator modules and their actions. Each modules is directed at a certain error but will also identify a number of other errors.

| Problem |  | SSR | NHHR | GIR | CGSR | PR |
| --- | --- | --- | --- | --- | --- | --- |
| Sequence error | ATG/Stop | X |  | X | X | X |
|  | Splice site | X |  | X | X | X |
| Pseudo-gene | ATG/Stop | X |  | X | X | X |
|  | Splice site | X |  | X | X | X |
|  | Similarity |  | X |  |  | X |
| Non-Homologous Hit |  | X | X |  |  | X |

The first step is the Short Sequence Remover (SSR) directed at short sequences. For this, a reference sequence is included in the preparation of the dataset. Sequences of proteins for which a structure has been resolved are preferred since these typically concern the final active protein, possibly following post-translational modifications such as propeptide cleavage. By default, sequences that have a length of 65% or less of the reference sequence are removed.

The second step is the Non-Homologous Hit Remover (NHHR) which removes outliers and is meant to remove non-homologous sequences. It constructs a HMMER profile and screens all sequences by hmmsearch. Sequences are removed according to the 3σ rule applied to total HMMER scores: Sequences with a score lower than the mean minus three standard deviations are removed, which, in a data set with normal distribution, corresponds with 0.3% of the sequences.

The third module in the default pipeline is the Gap Instigator Remover (GIR) which removes sequences that instigate large gaps in MSAs. By default, the GIR removes sequences that induce regions of 30 or more continuous gap columns, where a gap column is defined as a column that has 90% or more gaps, in order to circumvent problems with certain residues that may have become aligned to the insert. This default region size setting is based on the minimal intron size observed among most eukaryotes but may depend on the organism [34]. Sequences with additional N- and/or C-terminal subsequences will also be removed. Sequences are removed on a one-by-one basis (the sequence that instigates the largest gap first) and GIR is iterated, where iterations start with realignment.

The fourth module is the Continuous Gap Sequence Remover (CGSR). This removes sequences that show one or more instances of large continuous gaps in the MSA. Again default setting is at 30 columns whereas only gaps in columns with less than 50% gaps are considered. This is in order to allow subfamily specific subsequences. CGSR removes sequences in a threshold-controlled batch (See supplemental Document 1 for more detail) and is iterated. Note that not all sequences that lack a subsequence due to incorrect gene modelling will be detected. Residues that enclose the absent subsequence may align somewhere in the corresponding area, thereby splitting the region in two or more gap regions.

The last module in the default pipeline is the Pseudogene Remover (PR) which is identical to the NHHR module except that it is iterated. The names of the NHHR and PR modules are as such based on the intent of the modules. Although there is no clear-cut threshold that can discriminate between a non-homologous hit and a pseudogene, the first can be expected to show less similarity and be more disturbing. As such, NHHR is by default the first module and not iterated. PR is iterated since pseudogene identification is more delicate and works with a much improved MSA that corresponds with a different HMMER score distribution. As a result, PR is more sensitive than NHHR and by default the last module of the pipeline.

The third block is the recovery analysis since we envisage certain FH sequences will be inadvertently removed. Recovery analysis is performed correct overzealous sequence removal. The Seqrutinator output provides data that show the scrutiny in detail including some graphs that can assist in quality analysis. We performed a recovery analysis since there is no benchmark dataset available to quantitatively determine the performance of Seqrutinator.

### Design of the Performance Analysis

In order to develop automated sequence scrutiny, we generated the algorithms while testing their performance using superfamily datasets. Although the basic problems are straightforward, the implementation of any quantitative scrutiny is hampered by the fact that it will be difficult to use reliable cut-off thresholds and settings. The performance of a binary classifier such as Seqrutinator is usually determined in terms of precision and recall. NFHs are referred to as relevant instances, as real positives when detected and as false negatives when not detected. FHs detected as NFH are false positives. Unfortunately, determining error rates is not possible since no benchmark datasets are available. Hence, we need other methods to shed light on Seqrutinator performance. UniProtKB/Swiss-Prot (SwissProt) is, to the best of our knowledge, the most appropriate dataset for qualitative benchmarking of the pipeline. It contains a large number of sequences that come with biochemical and/or transcript evidence. However, it does also contain sequences that do not have any actual evidence for functionality. Hence, we can expect that some sequences will be NFH sequences.

We tested the pipeline on superfamilies with many homologues to allow for the statistical approach used in the outlier modules. We selected superfamilies involved in specialized metabolism since this provides a number of challenges. First, enzymes from specialized metabolism are under less functional constraint than enzymes from primary metabolism [35] by which datasets will show high sequence diversity. Then, specialized metabolism generates a plethora of chemically related compounds via parallel, diverging and converging pathways [35]. We envisage that promiscuous enzyme activity is often related to specialized metabolism and superfamily enzymes. The functional constraint will often act on (part of) the superfamily rather than on specific paralogues, which contributes to sequence variation. The complexity of the superfamilies we will use is so high that no reliable function annotation is at hand for many homologues. Note that enzyme promiscuity can result in incorrect sequence annotation. the last issue we foresee is that certain subfamilies are taxon-specific which may result in the inadvertent removal of FH sequences.

We tested the pipeline by simultaneously performing sequence mining for three functionally related and complex superfamilies in 16 complete plant proteomes. Cytochrome P450 (CYP) and UDP-Glycosyl Transferase (UGT) constitute 4.17% and 7.81%, respectively, of the known flavonoid metabolism in potato. The BAHD acyltransferase superfamily derives its name from its major enzymes: benzylalcohol O-acetyl transferase; anthocyanin O-hydroxycinnamoyl transferase; N-hydroxycinnamoyl anthranilate benzoyl transferase; and deacetylvindoline 4-O-acetyltransferase, constitutes a further 11.98%, of a total of 384 proteins that are classified in 59 Pfam domains (see Figure 2). As such, these three superfamilies form part of our major biological research interest which is to model flavonoid metabolism in potato (*Solanum tuberosum*). As such we need to functionally assign sequences of these three complex superfamilies. This can be performed by for instance Panther [36], which assigns at the subfamily level, rather than Pfam [9] which assigns at the superfamily level. Recently we developed HMMERCTTER [37], a software for the clustering and classification of protein superfamily sequences, which outperformed Panther in classifying three protein superfamilies. HMMERCTTER classification shows 100% precision and recall (100% P&R) but needs reliable sequence sets, i.e. sets that contain only a few NFH sequences.

**Figure 2:**
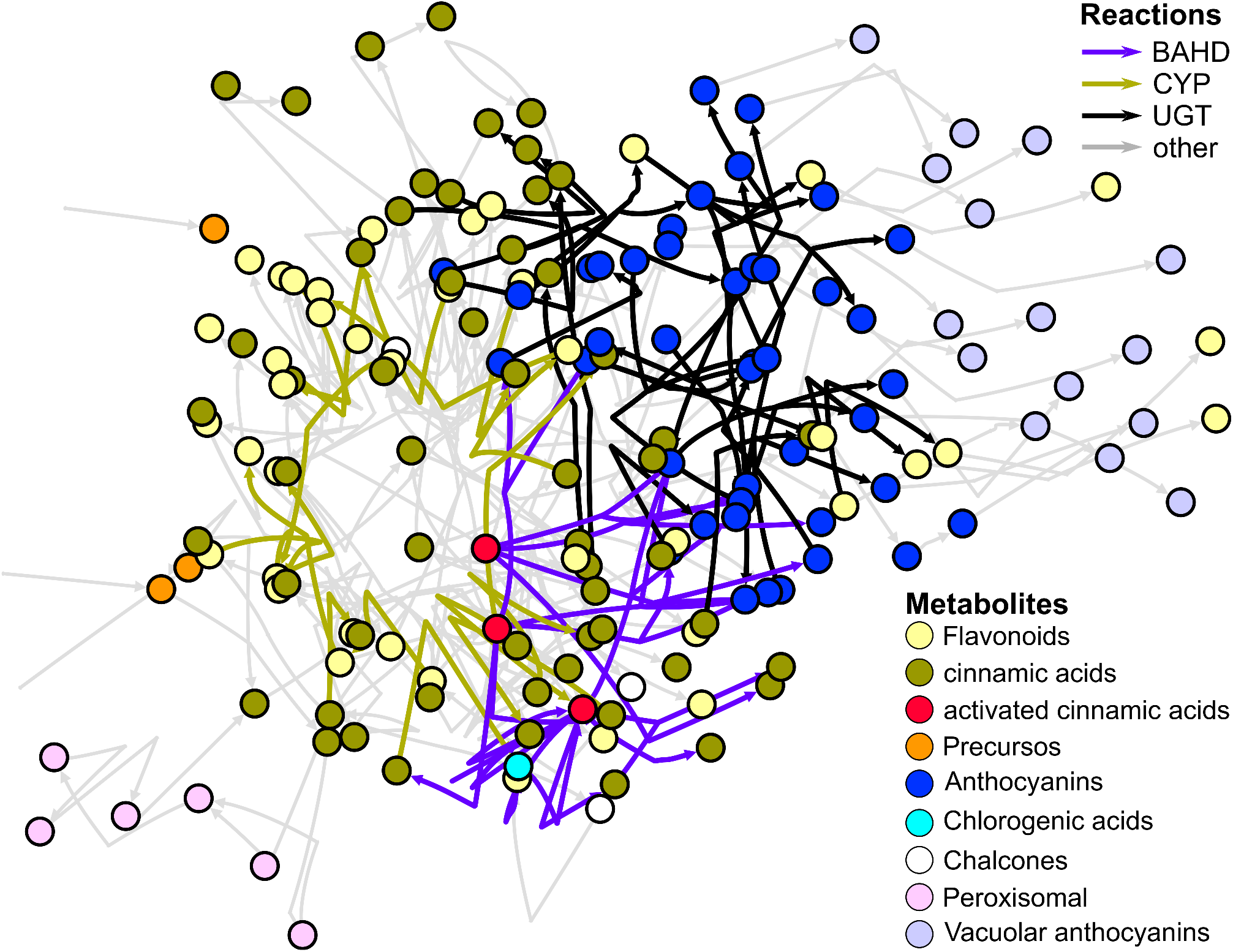
Cartoon Depicting the Phenylpropanoid/Flavonoid Biosynthesis Network in Potato. Three major superfamilies cover approximately 25% of the enzymatic reactions. The superfamily subnetworks show a partial overlap. The BAHD and UGT superfamilies are shifted towards end-products that are conjugated and transported to the vacuole, thereby changing the effective sink.

Automated sequence scrutiny of three superfamilies in 16 higher plants allows for a comparative analysis that will shed light on its performance. This is based on the hypothesis that performance on complete proteomes depends on the quality of the complete proteome rather than the superfamily that is analysed. Each of the applied algorithms should identify no or only a few NFH sequences in a high-quality sequence set such as the complete proteome from the model organism *Arabidopsis thaliana* (TAIR10). On the other hand, complete proteomes that have either been published only recently or cannot count on a large research community, are more likely to contain many NFH sequences. There are large differences between the number of sequences of the complete proteomes and although differences consist in the number of actual paralogues in different plants, we expect a certain level of convergence. For *A. thaliana* we included TAIR v6, besides the latest and supposedly superb set of TAIR v10. The actual superfamily analyses will be published elsewhere, here we report the numerical data from the sequence scrutiny to show the performance of the methods.

Finally, we considered how to detect false positives, i.e. inadvertently removed FH sequences. To do so we must understand the method and the biases of the initial datasets. For instance, a complete proteome from a single organism will have a different bias than the SwissProt sequence set or, for instance, all sequences from a single Pfam seed alignment.

A first minor concern is that of sequences that have been removed by the PR module because of a biased HMMER profile. Although HMMER profiles are weighted, they cannot account for large differences in clade distance. In a superfamily with various equidistant subfamilies and a single, more distant subfamily, sequences of the distant subfamily will show low scores in a hmmsearch and might have been removed by the PR. This problem is exacerbated by our approach of scrutinizing complete proteomes, which come with divergent MSAs and HMMER profiles with low precision and recall. These inadvertently removed sequences can be identified by cluster analyses of the combined sequences that were removed from the various individual sequence sets.

Another major concern is that of taxon-specific sequences. Complex superfamilies show a high rate of evolution by which taxon-specific sequences should be expected. Any type of mutation can result in a novel functional subfamily and as such functional sequences can have been removed by GIR, CGSR or PR. Cluster analysis is likely to fail when the sequence is taxon-specific, hence additional sequence mining may be required in order to enlarge the sequence space that is to be analysed.

## Results and Discussion

### Comparison Of The Pipeline On 19 Complete Proteomes Shows It Is Consistent

We subjected 19 sequence sets to sensitive HMMER searches with HMMER profiles for BAHD, CYP and UGT. This resulted in a total of three times 19 sequence sets representing the crude BAHDomes, CYPomes and UGTomes of the 16 plant species (Figure 3A), of which *A. thaliana* is represented with v6 and v10 of its complete proteome, as well as two SwissProt plant sequence sets (standard and with protein/ transcript evidence, from here on referred to as curated). All crude sequence sets were then prepared for and subjected to the Seqrutinator pipeline using default settings. The numbers of homologues after each module were recorded can be found in Supplemental File 1 and are resumed in Figure 3B. This shows similar patterns for the scrutiny and sequence removal on the three superfamilies were obtained, which suggests scrutiny performance is consistent.

**Figure 3:**
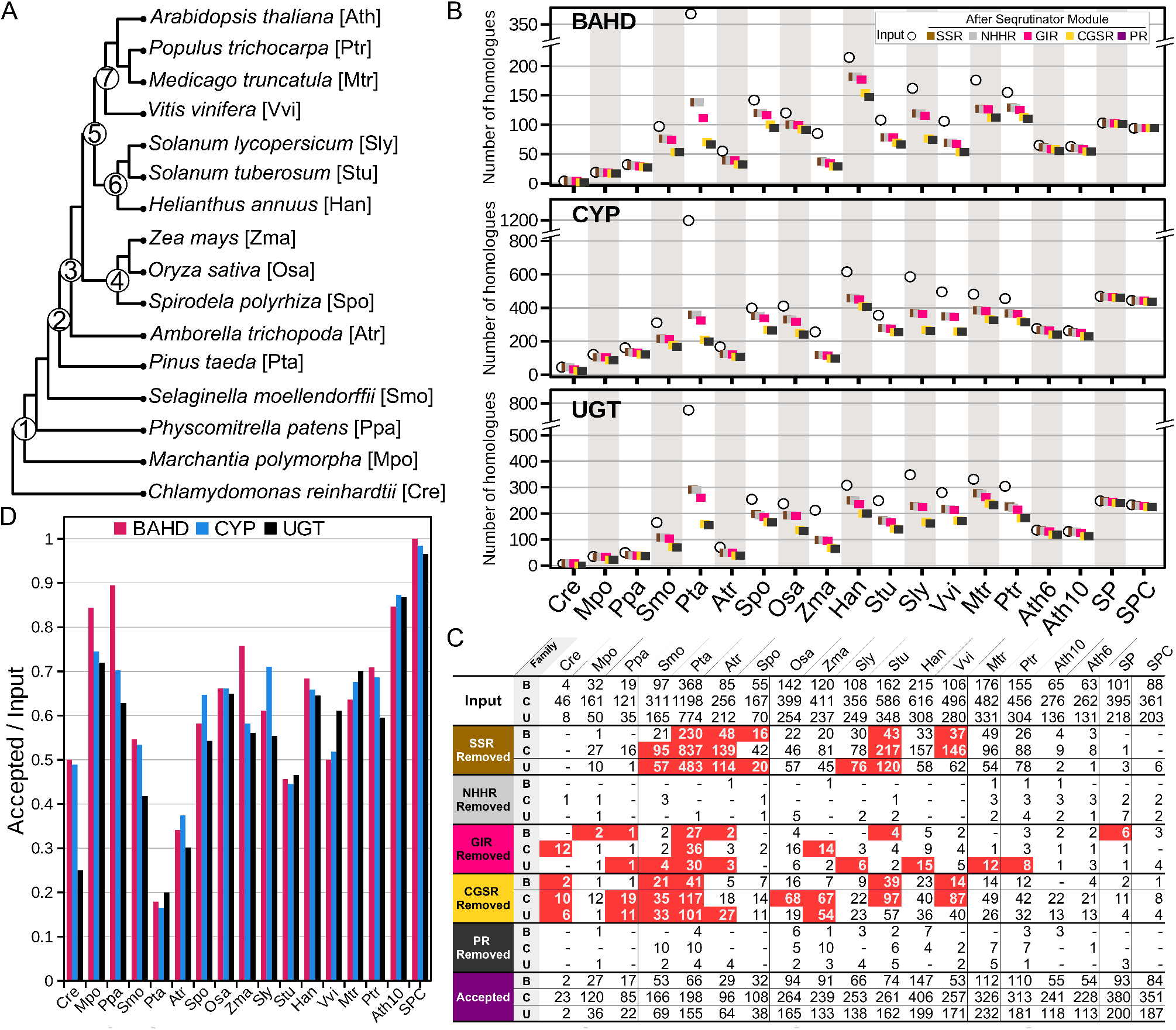
Seqrutinator Performance on 19 BAHDomes, CYPomes and UGTomes. **A:** Taxonomy of selected species. **B:** Numbers of BAHD, CYP and UGT homologues per species found and recovered after each step of the Seqrutinator pipeline. **C:** Number of Removed Sequences (default pipeline). Shown are the numbers of the initial and finally accepted sequences as well as the number of removed sequences per module and superfamily (B: BAHD, C: CYP and U: UGT). Red shading indicates a proportionally high number of NFH was removed (see also main text and Supplemental File 1, SSR, GIR and CGSR only). **D:** Seqrutinator performance for BAHDomes, CYPomes and UGTomes. Shown is the proportion of the numbers of finally accepted sequences and the numbers of initial sequences per species.

The table (Figure 3C) shows how many sequences were removed by each module from each of the three times 19 datasets. We normalized the number of removed sequences *n* (per module, superfamily and complete proteome) by expressing them proportionally to the number of finally accepted sequences *N*. This n/N ratio is a relative indication of how many sequences were removed and was used to compare the performance of the SSR, GIR and CGSR modules for each superfamily (the NHHR and the PR module remove too few sequences for meaningful comparison). Figure 3C highlights which modules removed more sequences than the average for which superfamilies. As hypothesized, relatively few sequences were removed from the *A. thaliana* datasets. No real difference was found comparing v6 with the more recent v10. Also, the *Marchantia polymorpha* (duckweed), *Oryza sativa* (rice), *H. annuus* (sunflower), *Medicago truncatula* (barrelclover) *and Populus trichocarpa* (black cottonwood poplar) datasets appear to have only a few NFHs. GIR removed six sequences from the SwissProt dataset but only three from the SwissProt with protein/transcript evidence dataset. *Pinus taeda* (loblolly pine), on the other hand, shows consistently many NFHs, detected by the SSR, GIR and CGSR modules from all three superfamily datasets. Note that this result, rather than for instance the detection of many outliers, corresponds with a low-quality proteome. Also, lycophyte *Selaginella moellendorffii, Amborella trichopoda* and *S. tuberosum* appear to have many NFHs removed by SSR, GIR and CGSR. Hence, it seems the performance of the modules is largely explained by the complete proteome dataset provided rather than by the superfamily. This is in correspondence with our hypothesis and seems to indicate that Seqrutinator has a good performance. On a side note, it seems that a number of genome consortia have applied sequence scrutinies towards partial sequences.

The most effective step is SSR, which indicates that many sequences, in particular from the complete proteome of *P. taeda*, are partial. Both versions of the *A. thaliana* complete proteome appear with a few partials (3, 8 and 1 for BAHD, CYP and UGT, respectively for v10). CGSR also removed many sequences which is explained by the relaxed setting of SSR (<65% of length reference sequence). GIR was the third most efficient module and removed, except for *P. taeda* and the algae *Chlamydomonas reinhardtii*, a few sequences per dataset. It appears to remove relatively many sequences from the UGT superfamily which suggests this case comes with false positives.

Figure 3D shows how many sequences are classified as FHs, relative to the initial number of sequences, for each species and each superfamily. Clearly, the complete proteome of *A. thaliana* appears as the best, with 85, 87 and 86% of each of the originally identified BAHD, CYP and UGT sequences classified as functional. On the other hand, the recently published complete proteome of *P. taeda* appears as very poor with a mere 18, 16 and 20% of the sequences classified as functional. The largest differences in scrutiny numbers amongst superfamilies are found for *Zea mays* (corn, 76, 58 and 56%) and *Solanum lycopersicum* (tomato, 61, 71 and 55%). However, this may be the result of differences in the initial number of homologues. For example, *S. lycopersicum* has relatively few, 356, CYP homologues of which 267 are considered functional (*S. tuberosum* has 586 homologues of which 253 are considered functional). *Z. mays* has relatively few, 120, BAHD homologues of which 91 are functional (*O. sativa* has 142 homologues of which 94 are considered as functional). Hence, although the numbers of removed sequences can differ, the final numbers of accepted homologues are similar which suggests that sequence scrutiny was consistent.

Most importantly, the SwissProt sequence set appears to have only a few sequences that were detected as NFH (4.6, 2.8 and 7.9% of the curated SwissProt dataset (see supplemental File 1)). This suggests the scrutiny was not overzealous but recovery analysis will have to show if these removed sequences are indeed not functional.

### Removal of NFH sequences Results in Improved MSA Quality

The ultimate goal of Seqrutinator is to obtain a sequence set that can be aligned with high fidelity. There are several methods that can be used to calculate the quality of an MSA. TCS [38] is one of the most accurate measures but, besides that it is computationally expensive, it does not provide highly discriminative scores, which makes it a difficult method for benchmarking. Hence, we sought alternative methods. A simple method is to look at the length of the MSA, as compared to the length of the mature protein. Now although this is a quantitative measure, MSA length does not accurately describe its quality since, for instance, a single large insert leads to a large MSA but does not necessarily result in either a good or bad MSA. As such, also the sum-of-pairs [39] is also not a very discriminative score.

A more sophisticated method is to determine the number of columns of trimmed MSAs. Since MSAs of complex superfamilies by definition have regions that are either not too reliable or specific to certain subfamilies (hence not truly homologous), MSAs are usually trimmed prior to phylogenetic reconstruction. Trimmers such as BMGE [32] or trimAl [33] remove columns with either high amounts of gaps or high entropy. As such, the length of an MSA following trimming conceptually reflects the number of reliable columns and can as such be used as a quality measure to compare MSAs of the same or, as in this case, similar datasets. Figure 4 shows the result of the scrutinies we performed. The number of reliable columns increases in almost all steps for all species and the three superfamilies. Besides that the MSAs clearly improve, typically with a factor 2 to 4, there appears to be a convergence of trimmed MSA length. Note that the nonseed plants scrutinies end with only a few sequences (Figure 3), by which the MSAs show low complexity and are typically larger following trimming (Figure 4). Another interesting detail is that, among a few others, the initial *P. taeda* MSAs of all three superfamilies appear completely unreliable. On the other end, the trimmed MSASs from the SwissProt and *A. thaliana* datasets increase in length. In general, the largest increases in trimmed MSA length occurs following the SSR and CGSR steps. This despite the fact that GIR removes sequences that induce large gap areas. This is likely related to the large number of sequences these modules removed.

Thus, Seqrutinator results in sequence sets that show significant improvement in quality, as demonstrated by the increase of the number of reliable columns of MSAs.

### Comparison of Seqrutinator Module Performance

The performance of the five scrutiny modules was analysed. First of all, we wondered how changing the cut-off threshold of the outlier modules affects the results. Rather than the default setting of 3σ (Means - 3 standard deviations), we applied 2.35σ, which corresponds with 95% inclusion according to a normal distribution. We performed analyses with different module orders with the major goal to determine if different modules detect the same NFHs. We tested pipe 4235 to see whether CGSR can replace SSR and at what cost. We tested pipe 134 to see the effect of omitting outlier removal and to test if GIR and CGSR detect outliers. Pfam scans with different cut-off thresholds were included as an external reference in order to shed light on performance in terms of precision and recall. Figure 6A shows the design and results of the analyses.

**Figure 5:**
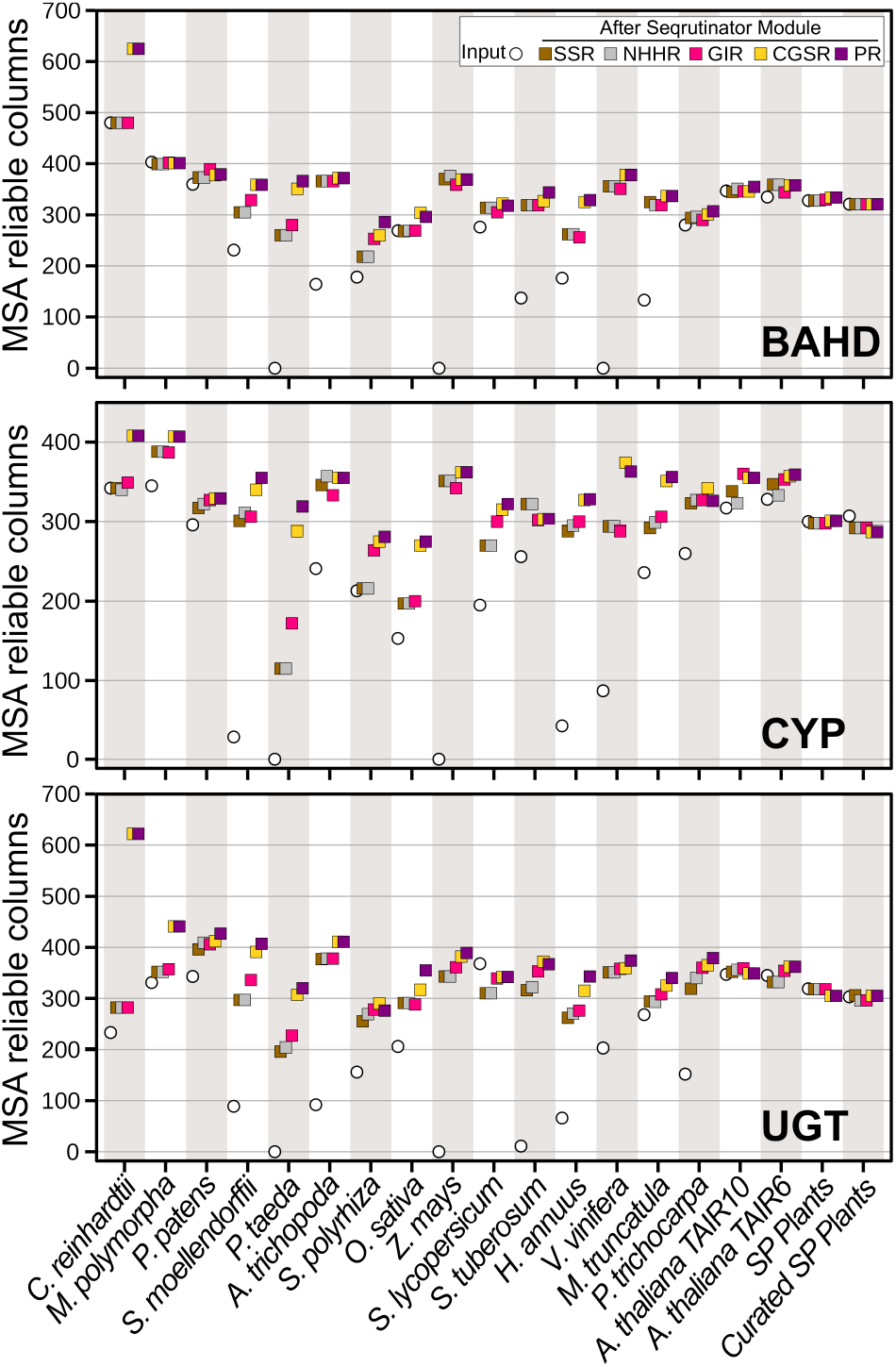
Number of Reliable Columns of MSAs After Each Step of the Seqrutinator Pipeline.

**Figure 6:**
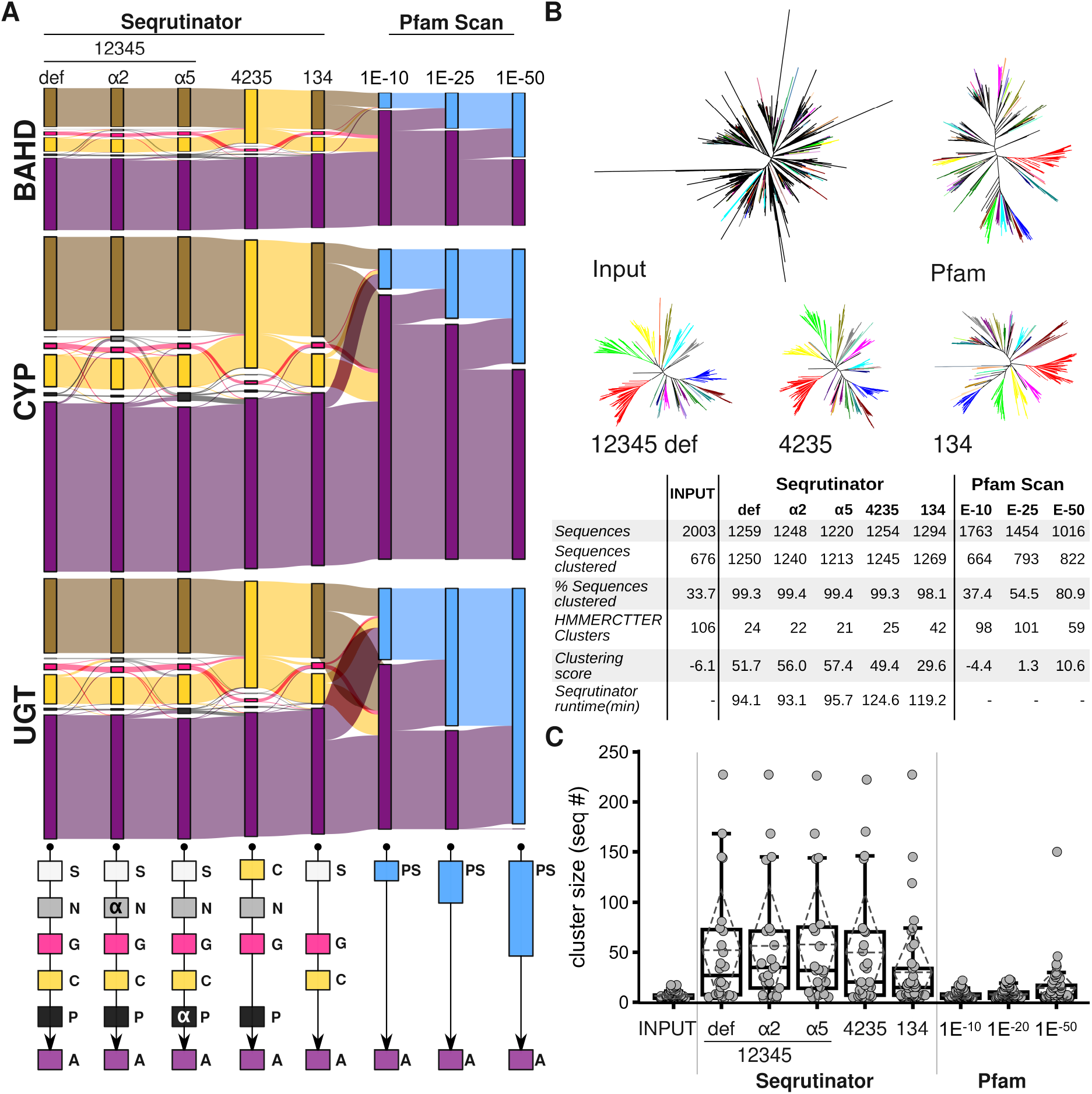
Seqrutinator is a Robust and Flexible Pipeline. **A:** Sequence fate in scrutinies with different pipelines. Top: Alluvial plot showing the fate of initial BAHD, CYP and UGT representatives (2003, 6782 and 3994 sequences respectively) in each of the applied scrutiny methods. Bottom: Schematic illustration of applied pipelines (S: SSR (1); N: NHHR (2); G: GIR (3) C: CGSR (4); P: PR (5); PS: Pfam Scan with cut-off thresholds as indicated; and A: accepted). The outlier removal cut-off was tested by comparing the default 3σ to the more strict 2.35σ cut-off in NHHR (α2) or PR (α5). **B:** HMMERCTTER clustering of BAHD sequence sets. Top: Cluster-wise coloured ML trees of five BAHD sequence-sets as indicated: Input: Partition of initial sequences; Pfam: Partition of sequences obtained with highly significant Pfam cutoff threshold (Expect value 1E-50); 12345 def: Partition of sequences accepted by default Seqrutinator pipeline; 4235 and 134: Partitions of sequences accepted by alternative Seqrutinator pipelines. Each cluster is assigned a different colour, black leaves are orphan (not clustered) sequences. Bottom: Numerical abstract of clustering analysis of all nine tested datasets. Shown are the total number of sequences, the number and percentage of clustered sequences, the number of clusters and the cluster scores, ((Clustered sequences-Orphans)/Total Sequences). **C:** Boxplots of cluster sizes of obtained HMMERCTTER partitions. The dotted lines show the mean and the standard deviation.

Alluvial plots show the fate of the sequences in all pipelines and Pfam scans. Interestingly we see most exchange combinations occur. For instance, in the 4235 pipe, CGSR not only takes care of all sequences SSR removes in the default pipeline, it also removes a number of sequences that are normally removed by GIR. This confirms the idea that many NFH sequences suffer from more than one of the initially described problems. The results with different outlier cut-offs suggest that particularly for PR, a more stringent screen will yield too many false positives.

The comparison with the Pfam scans also sheds light on performance. The original sequence sets were obtained with Pfam profiles but included all sequences with a score above HMMERs inclusion threshold. Pfam normally applies a more strict gathering threshold for each profile: a bitscore and corresponding E-value are set such that all sequences from the seed alignment are included. Note that application of the gathering threshold can result in missing homologues, as was the case for four BAHD sequences. We applied more stringent cut-offs in our analyses. Except for the UGTome, the most stringent cut-off of 1E-50 still accepts many partial sequences that are removed by either SSR or CGSR. A similar result is observed for GIR. On the other hand, many sequences that are not removed by Seqrutinator, hence accepted in the final dataset, are not accepted by Pfam using these stringent cut-offs. As such we conclude that Pfam scans suffer from poor precision and recall, which is a recurrent issue in clustering and classification and related to the fact that single profiles for complex superfamilies are used. The most conspicuous Pfam scan result was obtained when we applied 1E-50 to the UGT dataset. Only 12 sequences were accepted as FH, of which intriguingly one is normally removed by the SSR and another one by the CGSR module. The last detail cannot be observed in this particular alluvial plot due to the order of the datasets we applied in the figure.

Since function is conserved, we assume that functional families show mutual preservation where NFHs show lower conservation levels. HMMERCTTER clustering is a method in which superfamily sequences are clustered based on phylogeny and a HMMER score that is determined to include all sequences of a clade [37]. Only clusters with 100% P&R are accepted and as such HMMERCTTER partitions are conserved. If NFH sequences are less conserved clustering of a sequence set that lacks NFH sequences should therefore result in fewer but larger clusters. As such, we performed unguided HMMERCTTER clustering on the BAHD datasets and compared the resulting partitions (Figure 6B).

As expected, HMMERCTTER clustering of the crude dataset results in a very poor partition, with more orphans than clustered sequences as shown by the negative cluster score (*(Clustered sequences-Orphans)/Final Sequences*). It should be noted that HMMERCTTER clustering uses score rather than E-value by which many partial sequences will end up as orphan sequence. The Pfam 1E-50 tree shows much fewer orphans and has a much better cluster score. The best cluster score (57.4, see Table in Figure 6B) is obtained with the dataset that results from the 1234a5 pipeline in which a substantial number of low-scoring outliers has been removed. The default pipeline has cluster score of 51.7 which reflects a trade-off between a larger number of accepted sequences and a slightly lower number of clusters. As compared to the default pipeline, the 4235 pipeline has a slightly lower performance which, combined with its substantially longer runtime, shows the default pipeline is preferred. Whether a more strict cut-off should be applied for the PR module depends on whether high precision or high sensitivity is preferred. Not applying any outlier module (pipe 134) results however in a substantially reduced cluster score of 29.6, even though only a few outliers are detected.

Figure 6B shows that, based on the above-made assumption, the application of Seqrutinator in superfamily sequence mining results in largely improved datasets. This is also reflected by the distribution of cluster size (Figure 6C). Although Pfam scan with a very strict cut-off threshold of 1E-50 shows an increase in cluster-size, it has very few large clusters (>50 sequences per cluster), as compared to the Seqrutinator-derived datasets.

### Recovery Screen of SwissProt BAHDome, CYPome and UGTome

We performed a recovery screen directed at the detection of false positives, i.e. sequences of FHs that were inadvertently removed by the automated scrutiny. The recovery screens consisted largely of a number of alignment analyses directed at understanding why certain sequences were removed. In general, this can be performed in order to recover inadvertently removed sequences. In this study, the major goal was to show if scrutiny was not overzealous. Hence first, we applied recovery screens to the SwissProt-based datasets that served as a positive control. Table 2 summarizes the results and shows performance in terms of false positives. Supplemental document 2 contains an elaborate description of the complete recovery screen including the analysis of the three 16-proteome sequence scrutinies.

**Table 2:**
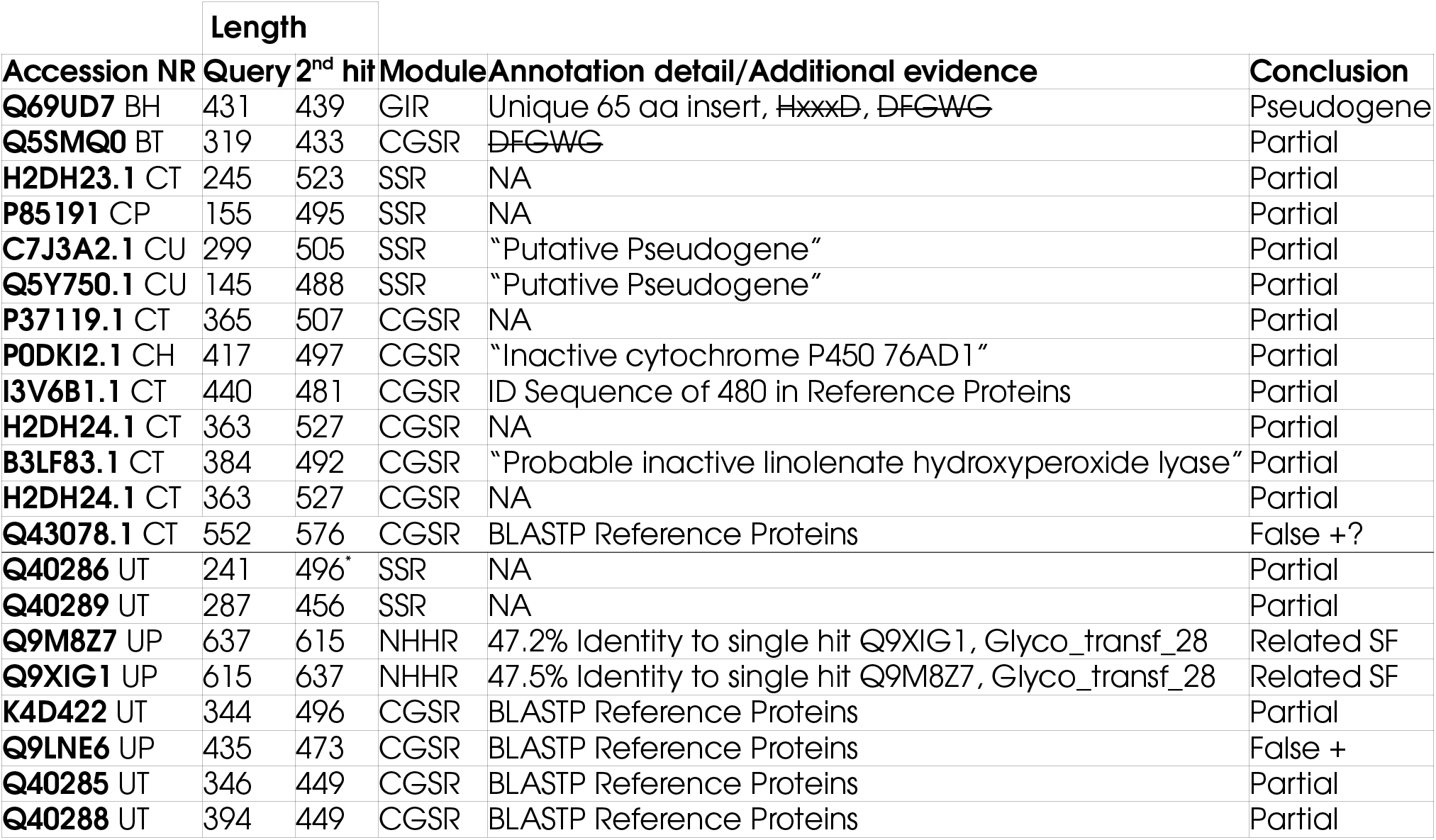
Recovery Analysis SwissProt Datasets. B, C and U indicate BAHD, CYP and UGT case respectively. P and T indicate sequences with protein or transcript evidence, H means inferred from homology, U uncertain. * Third hit since second hit, K4D422, was not representative. False + means the sequence was incorrectly identified as NFH. NA means Not Applicable.

Not a single sequence was removed from the curated BAHD SwissProt set, two sequences were removed from the non-curated SwissProt sequence set. Despite that its total length of 431 residues is similar to what is found for BAHD sequences, Q69UD7 was removed by GIR because of a unique ~65-residue long insert. Furthermore, two of the key motifs defining the BAHD superfamily (HxxxD and DFGWG) are missing in this entry. Then, a BLAST search vs. the SwissProt database shows that the second hit (Q0DKA5) has a mere 38.8% of identity with the query. Together, these results suggest that Q69UD7 is a pseudogene. Q5SMQ0 appears to be a partial sequence that lacks a C-terminal fragment with the DFGWG motif.

All but one SwissProt sequences that were removed from the initial CYP homologue set were partial and as such correctly removed (See Table 2). Most BLAST alignments we performed with these SwissProt sequences against SwissProt showed the absence of either or both the N- and the C-terminus. Only Q43078.1 showed an internal gap, as demonstrated by aligning its sequence to the final MSA using MAFFT add. BLASTP against SwissProt, however, showed no large gap suggesting this sequence may represent an FH and a first false positive. None of the sequences had evidence at the protein level and some sequences had annotations that suggested they are NFHs. Two sequences were removed from the non-curated SwissProt dataset by CGSR. The curated dataset had two additional sequences removed, i.e. sequences that were not removed from the non-curated dataset. This indicates that although the procedure is fully objective, its performance depends on the data.

We also performed an analysis of the sequence scrutiny obtained with NHHR applying a more strict cut-off threshold. This did remove a number of FHs. Orthologues of these SwissProt sequences were sometimes removed from a complete proteome by the PR module with the default setting. Hence, the strict NHHR scrutiny of the well-annotated SwissProt set can be seen as predictive for the performance on other datasets as is described in detail in Supplemental document 2. This also shows how some automatically generated graphs can assist in the recovery analysis.

In the case of the UGTome, eight sequences with either protein or transcript evidence were removed. The two sequences removed by NHHR show over 47% mutual identity. Pfam scan indicates these are enzymes from the related protein family of Glycosyl Transferase 28 (GT28, PF03033). Interestingly, similar instances of GT28 sequence homologues were removed by NHHR (6), GIR (12), CGSR (33), and PR (10) from the 16 proteomes. A more profound analysis is described in Supplemental Document 2 and shows these are indeed not UGTs. Not a single GT28 sequence was retained as UGT by Seqrutinator. This indicates that Seqrutinator is also sensitive albeit that NHHR did not detect all GT28 sequences. Since PR at the same threshold did identify all GT28 sequences that were missed by NHHR, GIR and CGSR, we conclude that this is caused by the rather poor initial MSA. Note that these results also explain the relatively high number of sequences that GIR removed from particularly the UGT datasets as was shown earlier (Figures 3B and 3C).

Four putative UGT sequences were removed by CGSR. When aligned to the accepted SwissProt sequences, all appear to lack N-terminal sequence. This was further verified for three of the four sequences by BLASTP analysis against the Reference Protein database. The fourth sequence, Q9LNE6, appears to have a number of homologues that align well as of from the first residues, which suggests it concerns another possible false positive that may correspond to an FH. Indeed, this sequence was supported by evidence at the protein level.

Resumed, the analyses of both the SwissProt sequence sets and the 16-proteome datasets (See Supplemental document 2) show that Seqrutinator detects mostly real NFH but also that in certain occasions some FHs may be removed. Perhaps more importantly, the recovery analysis shows that, since Seqrutinator generates a dataset with a high quality MSA, it allows to perform a fast recovery analysis and therewith correct eventual overzealous sequence removal. Alignment to an existing MSA is straightforward using MAFFT add. Other recommended analyses are CD Hit cluster analysis and Pfam scan (See examples of their use in Supplemental Document 2).

### The Effect of the Data on the Performance

Next we analysed if the size of the offered dataset affects performance. Outlier detection, here performed with the 3σ rule, is mostly applied for datasets that show a normal distribution, which is severely affected by the dataset size. The analysis was also instigated by differences in the behaviour of certain sequences present in both the SwissProt and complete proteome datasets. In all three studied cases, the SwissProt sequence sets were somewhat smaller in size than the complete proteome-derived datasets. However, the SwissProt datasets do show different variations of subfamily homologues, which may explain a number of the obtained results. As such, we tested Seqrutinator’s performance with differently-sized, randomly-generated datasets.

We used the BAHD dataset to create randomized sequence sets, using four different size conditions: V(ariable) (datasets emulating the sizes of the datasets generated by different proteomes, ranging from 4 to 368 sequences), S(mall) (40 x ~50 sequences, each subset representing ~2.5% of the total sequences), M(edium) (20 x ~99 sequences representing ~5% of the total sequences) and L(arge) (10x~199 sequences, representing ~10% of the total sequences). Ten random datasets were generated for each condition. Additionally, we combined all the sequences in a single dataset. All of these datasets were scrutinized with the default pipeline (12345) and a pipeline lacking the outlier removers (134). Due to the dataset size of the single combined dataset, this dataset was scrutinized using FAMSA rather than MAFFT G-INS-i as alignment method. The datasets as originally prepared were scrutinised using both pipelines using either FAMSA or MAFFT G-INS-i). The design is sketched in Figure 6A.

**Figure 6:**
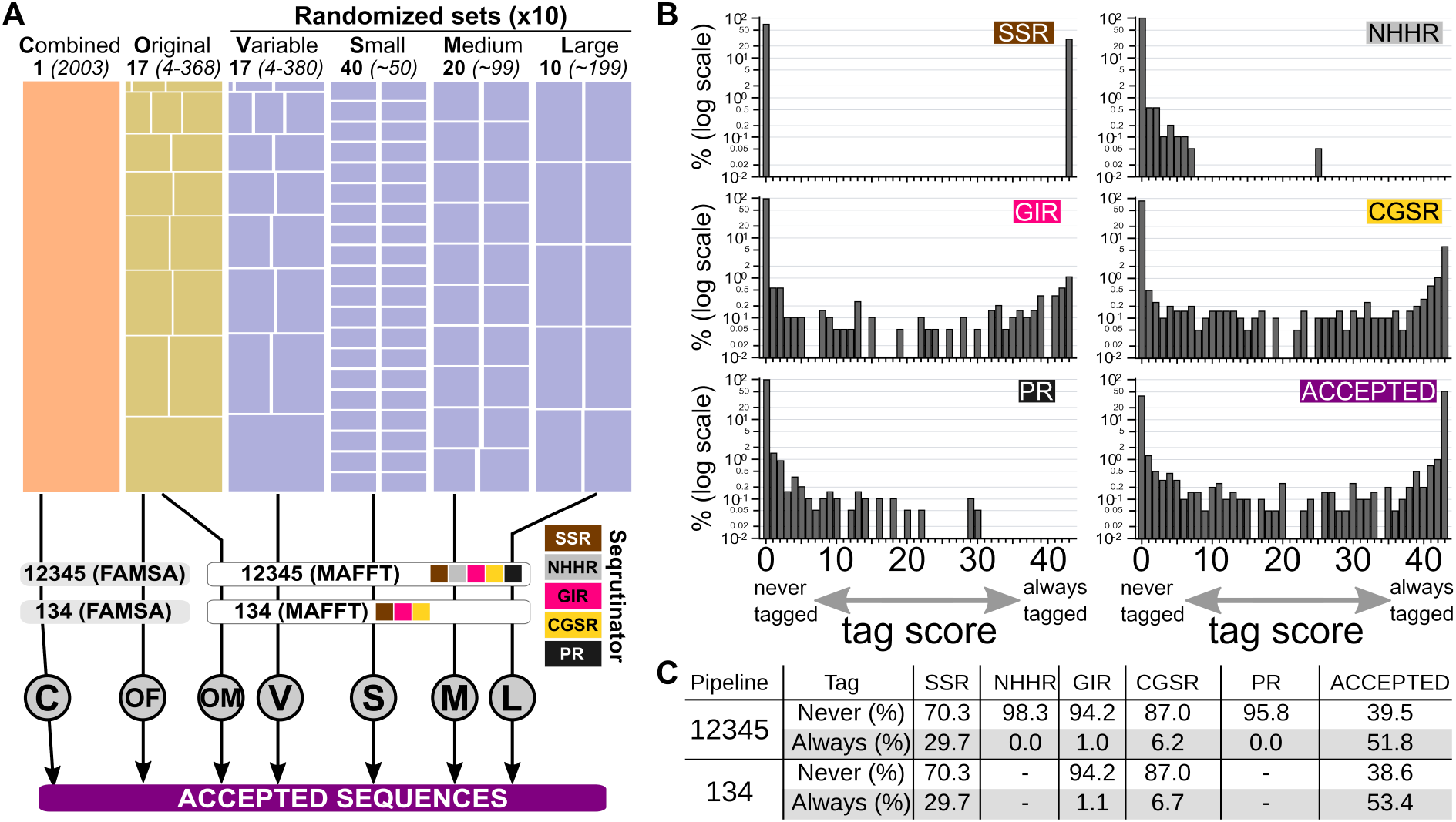
The Effect of Size on Seqrutinator Performance. **A: Design of Analysis.** Randomized datasets were compared with the original and the combined dataset. Four types of randomized datasets were made, each representing different subset sizes as indicated by the number of subsets (in bold) with the number of sequences per subset in brackets. Ten randomized sets were generated for each type and analysed using different scrutiny strategies as indicated. FAMSA was used in order to allow the analysis of the combined (C) dataset and used also for the original dataset (OM and OF; Original MAFFT and Original FAMSA). **B: Sequence Fate in Default scrutiny.** The tag score is the number of times a sequence was either removed by each module or finally accepted. The histograms show the incidence of the tag scores. **C: Final Sequence Fate.** Numbers of always rejected and always accepted for the 12345 and the 134 pipelines.

We first quantified the number of sequences tagged by each module for each dataset (Supplemental Figure 1). SSR showed no differences since it simply removes all sequences below a certain length determined by the reference sequence. The PR module removes more sequences from the large datasets. All other modules did show differences but no trend indicates that the size of the set affects the modules’ performance. The fact that NHHR does not show the same trend as PR is explained by the higher complexity of the MSAs made of sets with many more NFHs. Either way, NHHR and PR remove only a few homologues and present coefficients of variation (CV) of over 10% in all the randomized datasets. GIR removes at least 43 sequences with an average of 74 ± 4.05 per complete dataset. It has a CV of over 5% for both the 12345 and the 134 pipelines applied to variable and medium-size randomized sets. CGSR removes more sequences, ranging from 194 to 215 but shows low CVs, which suggests it is more reliable. Hence, size appears to affect only the PR module and the remaining differences are likely better explained by random variation.

As a next question, we wondered if the same sequences are removed and to what extent. Hence, for both the default and the 134 pipelines we determined how often each sequence was removed by each module, thereby generating a tag score for each sequence. Figure 6B shows the incidence per tag score for each module as part of the default pipeline, where 6C summarizes the total numbers for both pipelines. Clearly, by far the largest part of the sequences was either always (tag score Accepted = 0) or never (tag score Accepted = 43) removed, leaving only a few so-called twilight zone sequences with a fate that depends on the dataset. Only the CGSR module shows more than 5% of twilight zone sequences. Both the NHHR and the PR module data show a skew towards never removed.

We also wondered to which extent the same sequences are removed by different modules from different datasets. Here it concerns a total of 227 sequences with tag scores ranging from 1 to 42, removed by NHHR, GIR, CGSR or PR. These 227 remaining sequences were analysed for cross-module, cross-pipeline fate. It appears that a substantial part of these sequences can be removed by more than a single module, depending on the dataset (Supplemental Figure 2). About two-thirds of the sequences, 174 and 160 for pipes 12345 and 134, respectively, are sometimes accepted and sometimes rejected. Hence, overall, the performance of Seqrutinator depends only to a small amount on the exact dataset.

The above analyses also allow for additional analysis of the performance in terms of precision. We made the assumption that sequences that are always removed are real NFHs (RNFHs) whereas sequences that are never removed are real FHs (RFHs), as such creating what may be referred to as a silver standard. The remaining twilight zone sequences are, for this analysis, considered Putative Functional Homologues (PFH). We aligned the RFH sequences and then aligned, in two separate analyses the PFH and RNFH sequences to the obtained MSA. The two resulting MSAs were used to reconstruct phylogenies which were subjected to HMMERCTTER clustering with orphan and outlier removal (HMMERCTTER-OOR). This optional module detects and removes orphans (here defined as sequences that do not cluster due to the minimal cluster size-setting of N = 4) and putative outliers with the main goal of increasing cluster size. We then determined which of these orphans and outliers are NFHs or PFHs. Table 3 summarizes the results. The largest effect of including either NFHs or PFHs is that many sequences, irrespective of whether they represent a RFH, PFH or NFH, are detected as orphan. This is explained by the deterioration of the MSA and resulting tree. Both PFH and NFH but not RFH constitute a small number of outliers. Also, they represent proportionally more orphans than the RFHs. This suggests that at least part of what we considered NFH and PFH is indeed an NFH. This is supported by the fact that adding the sequences results in a substantial decrease of the number of reliable columns in the MSAs.

**Table 3:** HMMERCTTER-OOR Clustering of Non- and Putative Functional Homologues. Trees for the Real Functional Homologues (RFH) alone as well as combined with Putative Functional Homologues (R+PFH) and Non-Functional Homologues (R+NFH), respectively, were clustered with HMMERCTTER-OOR in order to identify the numbers of clustered, orphaned and outlier sequences. BMGE indicates the number of columns that were accepted as informative by BMGE.

|  | BMGE | Sequences | Total | Clustered | Outlier | Orphan | % Outlier | % Orphan |
| --- | --- | --- | --- | --- | --- | --- | --- | --- |
| **RFH Tree** | 133 | RFH | 1038 | 1033 | 0 | 5 | 0,00 | 0.48 |
| **R+PFH Tree** | 85 | RFH | 1038 | 1002 | 0 | 36 | 0,00 | 3.47 |
|  |  | PFH | 174 | 161 | 6 | 7 | 3.45 | 4.02 |
| **R+NFH Tree** | 77 | RFH | 1038 | 689 | 0 | 349 | 0,00 | 33.62 |
|  |  | NFH | 196 | 91 | 9 | 96 | 4.59 | 48.98 |

## Conclusion

We have made a flexible pipeline for the stepwise scrutiny of superfamily protein sequence sets with the objective of obtaining comprehensive sets that lack NFH sequences. Large, high quality MSAs of over 2000 FH sequences were obtained for three complex superfamilies. The MSAs have high quality, especially as compared with MSAs prior sequence scrutiny. Clustering analysis confirmed that the datasets improved significantly, yielding phylogenetic trees with cluster scores of over 50 and only very few orphans. Most clades, whether with or without SwissProt representative that may indicate function, have rather complete or at least broad taxonomic contributions. Although there is no benchmark set for sequence scrutiny, we show Seqrutinator has high precision and recall. Analysis of the SwissProt sequences showed that only a few sequences yield false positives, suggesting Seqrutinator has a high precision. In the scrutiny of the UGTomes, Seqrutinator correctly detected all GT28 sequences, which form a homologous superfamily, suggesting Seqrutinator has a high sensitivity or recall. Additional analysis in which we compared Seqrutinator to Pfam scan with different cut-offs, corroborated Seqrutinator has high precision and recall. The tool is flexible with five different scrutiny modules that also contribute to its robustness. The performance depends slightly on the size and complexity of the initial dataset. The method was designed for complex eukaryotic superfamilies but can be applied to intermediately complex and even simple superfamilies without any problem. The major exception is that it cannot deal with superfamilies with different domain architectures. Sequences from distant or taxon specific subfamilies may be removed inadvertently but the binary character of Seqrutinator allows for a simple recovery analysis.

## Materials and Methods

### Initial Datamining

The initial sequence mining was performed by MUFASA (See Supplemental Document 1) that applies hmmsearch from HMMER [40] using Pfam profiles PF02458 for BAHD [41], PF00067 for CYP [42]; and PF00201 for UGT [43] using HMMERs inclusion threshold as inclusion cut-off. Searches were performed in batch using the MUFASA script (See supplemental document 1) and the 16 plant species sequence sets obtained from Phytozome v12.1.6 [44], TAIR v6 that was obtained from TAIR [45], and the SwissProt datasets that were from Uniprot [46, 47]. Sequences for structures, PDB identifiers 4G0B (BAHD [48]), 5YLW (CYP [49]) and 3HBF (UGT [50]), were identified as top scoring sequence with the respective superfamily Pfam profiles.

### Sequence Alignment and other biocomputational analyses

All MSAs were performed using MAFFT-G-INS-i [51], except when indicated that FAMSA [25] was used. MSAs were analysed and shown using Aliview [52] or MSAviewer [53] at NCBI [54]. Trimming in Seqrutinator was performed using BMGE [32] using standard gap settings, BLOSUM62 and an entropy cut-off *h* of 0.8. Due to the poor quality of some datasets (e.g., dataset before Seqrutinator), which resulted in low or none reliable columns with BMGE, all datasets for phylogeny were first trimmed with trimAl with -gappyout settings followed by BMGE. CD Hit clustering [55] was performed at the CD Hit suite [56] at an identity cut-off of 0.3. BLAST [57, 58] analysis was performed at NCBI [59] against the database as indicated. Pfam scans were performed at EBI [60] using Pfam’s gathering threshld for cut-off or locally if and with Expect values indicated. Dotplots [61]were performed at the SIB [62]. Pyhylogenies were reconstructed by FastTree [19], using the WAG model and optimized Gamma20 likelihood, and drawn by Dendroscope [63]. Local alignments were performed with LALIGN/PLALIGN [64] at the UVA [65]. Alluvial diagrams were generated using RAWGraphs [66]. Boxplots and histograms were prepared with Plotly (Plotly Technologies Inc. Collaborative data science. Montréal, QC, 2015 [67]).

### Seqrutinator

A full description of Seqrutinator can be found in Supplemental document 1.

## Acknowledgements

AA and NS are CONICET fellows, AtH and VF are CONICET Career investigators. Funding was from ANPCyT, project PICT2017-1310.

## Cited References, Information Resources and webtools

1. Villarreal F, Stocchi N, ten Have A. Functional Classification and Characterization of the Fungal Glycoside Hydrolase 28 Protein Family. J Fungi. 2022;8:217. doi:10.3390/jof8030217.

2. Bondino HG, Valle EM, ten Have A. Evolution and functional diversification of the small heat shock protein/α-crystallin family in higher plants. Planta. 2012;235:1299–313. doi:10.1007/s00425-011-1575-9.

3. Bustamante JP, Radusky L, Boechi L, Estrin DA, ten Have A, Martí MA. Evolutionary and Functional Relationships in the Truncated Hemoglobin Family. PLoS Comput Biol. 2016;12:e1004701. doi:10.1371/journal.pcbi.1004701.

4. Valiñas MA, Have A ten, Andreu AB. Identification of the functions of 4-coumarate-CoA ligase/ acyl-CoA synthetase paralogs in potato. bioRxiv. 2021;:2021.07.06.451337. doi:10.1101/2021.07.06.451337.

5. Revuelta M V., van Kan JAL, Kay J, ten Have A. Extensive Expansion of A1 Family Aspartic Proteinases in Fungi Revealed by Evolutionary Analyses of 107 Complete Eukaryotic Proteomes. Genome Biol Evol. 2014;6:1480–94. doi:10.1093/gbe/evu110.

6. Kumar K, Mhetre A, Ratnaparkhi GS, Kamat SS. A Superfamily-wide Activity Atlas of Serine Hydrolases in Drosophila melanogaster. Biochemistry. 2021;60:1312–24. doi:10.1021/ACS.BIOCHEM.1C00171.

7. Spence MA, Mortimer MD, Buckle AM, Minh BQ, Jackson CJ. A Comprehensive Phylogenetic Analysis of the Serpin Superfamily. Mol Biol Evol. 2021;38:2915–29. doi:10.1093/MOLBEV/MSAB081.

8. Lin LM, Guo HY, Song X, Zhang DD, Long YH, Xing Z Bin. Adaptive Evolution of Chalcone Isomerase Superfamily in Fagaceae. Biochem Genet. 2021;59:491–505. doi:10.1007/S10528-020-10012-Z.

9. Finn RD, Bateman A, Clements J, Coggill P, Eberhardt RY, Eddy SR, et al. Pfam: The protein families database. Nucleic Acids Research. 2014;42:D222–30. doi:10.1093/nar/gkt1223.

10. Haft DH, Selengut JD, White O. The TIGRFAMs database of protein families. Nucleic Acids Research. 2003;31:371–3. doi:10.1093/nar/gkg128.

11. Orts F, ten Have A. Structure-function analysis of Sedolisins: Evolution of tripeptidyl peptidase and endopeptidase subfamilies in fungi. BMC Bioinformatics. 2018;19:464. doi:10.1186/s12859-018-2404-y.

12. Gough J, Karplus K, Hughey R, Chothia C. Assignment of homology to genome sequences using a library of hidden Markov models that represent all proteins of known structure. J Mol Biol. 2001;313:903–19. doi:10.1006/jmbi.2001.5080.

13. Simonetti FL, Teppa E, Chernomoretz A, Nielsen M, Marino Buslje C. MISTIC: Mutual information server to infer coevolution. Nucleic Acids Res. 2013;41 Web Server issue:W8–14. doi:10.1093/nar/gkt427.

14. Mazin P V, Gelfand MS, Mironov AA, Rakhmaninova AB, Rubinov AR, Russell RB, et al. An automated stochastic approach to the identification of the protein specificity determinants and functional subfamilies. Algorithms Mol Biol. 2010;5:29. doi:10.1186/1748-7188-5-29.

15. Wilkins A, Erdin S, Lua R, Lichtarge O. Evolutionary trace for prediction and redesign of protein functional sites. Methods Mol Biol. 2012;819:29–42. doi:10.1007/978-1-61779-465-0_3.

16. Chagoyen M, García-Martín JA, Pazos F. Practical analysis of specificity-determining residues in protein families. Brief Bioinform. 2016;17:255–61. doi:10.1093/BIB/BBV045.

17. Guindon S, Dufayard J-F, Lefort V, Anisimova M, Hordijk W, Gascuel O. New Algorithms and Methods to Estimate Maximum-Likelihood Phylogenies: Assessing the Performance of PhyML 3.0. Syst Biol. 2010;59:307–21. doi:10.1093/sysbio/syq010.

18. Stamatakis A. RAxML version 8: a tool for phylogenetic analysis and postanalysis of large phylogenies. Oxford University Press; 2014. doi:10.1093/bioinformatics/btu033.

19. Price MN, Dehal PS, Arkin AP. FastTree 2 - Approximately maximum-likelihood trees for large alignments. PLoS One. 2010;5:e9490. doi:10.1371/journal.pone.0009490.

20. Ronquist F, Huelsenbeck JP. MrBayes 3: Bayesian phylogenetic inference under mixed models. Bioinformatics. 2003;19:1572–4. doi:10.1093/bioinformatics/btg180.

21. Notredame C, Higgins DG, Heringa J. T-coffee: a novel method for fast and accurate multiple sequence alignment 1 1Edited by J. Thornton. J Mol Biol. 2000;302:205–17. doi:10.1006/jmbi.2000.4042.

22. Katoh K, Rozewicki J, Yamada KD. MAFFT online service: Multiple sequence alignment, interactive sequence choice and visualization. Brief Bioinform. 2018;20:1160–6. doi:10.1093/bib/bbx108.

23. Löytynoja A, Vilella AJ, Goldman N. Accurate extension of multiple sequence alignments using a phylogeny-aware graph algorithm. Bioinformatics. 2012;28:1684–91. doi:10.1093/BIOINFORMATICS/BTS198.

24. Szalkowski AM. Fast and robust multiple sequence alignment with phylogeny-aware gap placement. BMC Bioinformatics. 2012;13:1–11. doi:10.1186/1471-2105-13-129/TABLES/3.

25. Deorowicz S, Debudaj-Grabysz A, Gudys A. FAMSA: Fast and accurate multiple sequence alignment of huge protein families. Sci Rep. 2016;6:1–13. doi:10.1038/srep33964.

26. Shen C, Zaharias P, Warnow T. MAGUS+eHMMs: improved multiple sequence alignment accuracy for fragmentary sequences. Bioinformatics. 2022;38:918–24. doi:10.1093/BIOINFORMATICS/BTAB788.

27. Tumescheit C, Firth AE, Brown K. CIAlign: A highly customisable command line tool to clean, interpret and visualise multiple sequence alignments. PeerJ. 2022; 10:e12983. doi:10.7717/PEERJ.12983.

28. Chiner-Oms A, González-Candelas F. EvalMSA: A program to evaluate multiple sequence alignments and detect outliers. Evol Bioinforma. 2016;12:277–84. doi:10.4137/EBO.S40583.

29. Mendoza MLZ, Nygaard S, Da Fonseca RR. DivA: detection of non-homologous and very divergent regions in protein sequence alignments. BMC Res Notes. 2014;7. doi:10.1186/1756-0500-7-806.

30. Jehl P, Sievers F, Higgins DG. OD-seq: outlier detection in multiple sequence alignments. BMC Bioinformatics. 2015;16. doi:10.1186/S12859-015-0702-1.

31. Maldonado E, Antunes A. LMAP_S: Lightweight Multigene Alignment and Phylogeny eStimation. BMC Bioinformatics. 2019;20. doi:10.1186/s12859-019-3292-5.

32. Criscuolo A, Gribaldo S. BMGE (Block Mapping and Gathering with Entropy): A new software for selection of phylogenetic informative regions from multiple sequence alignments. BMC Evol Biol. 2010;10:210. doi:10.1186/1471-2148-10-210.

33. Capella-Gutiérrez S, Silla-Martínez JM, Gabaldón T. trimAl: a tool for automated alignment trimming in large-scale phylogenetic analyses. Bioinformatics. 2009;25:1972–3. doi:10.1093/bioinformatics/btp348.

34. Hong X, Scofield DG, Lynch M. Intron size, abundance, and distribution within untranslated regions of genes. Mol Biol Evol. 2006;23:2392–404. doi:10.1093/molbev/msl111.

35. Rieseberg TP, Dadras A, Fürst-Jansen JMR, Dhabalia Ashok A, Darienko T, de Vries S, et al. Crossroads in the evolution of plant specialized metabolism. Semin Cell Dev Biol. 2022. doi:10.1016/J.SEMCDB.2022.03.004.

36. Mi H, Muruganujan A, Thomas PD. PANTHER in 2013: modeling the evolution of gene function, and other gene attributes, in the context of phylogenetic trees. Nucleic Acids Res. 2013;41:D377–86. doi:10.1093/nar/gks1118.

37. Pagnuco IA, Revuelta MV, Bondino HG, Brun M, Ten Have A. HMMER cut-off threshold tool (HMMERCTTER): Supervised classification of superfamily protein sequences with a reliable cut-off threshold. PLoS One. 2018;13.

38. Chang J-M, Di?Tommaso P, Lefort V, Gascuel O, Notredame C. TCS: a web server for multiple sequence alignment evaluation and phylogenetic reconstruction: Figure 1. Nucleic Acids Res. 2015;43:W3–6. doi:10.1093/nar/gkv310.

39. Carrillo H, Lipman D. The Multiple Sequence Alignment Problem in Biology. http://dx.doi.org/101137/0148063. 2006;48:1073–82. doi:10.1137/0148063.

40. Eddy SR. A NEW GENERATION OF HOMOLOGY SEARCH TOOLS BASED ON PROBABILISTIC INFERENCE. In: Genome Informatics 2009. PUBLISHED BY IMPERIAL COLLEGE PRESS AND DISTRIBUTED BY WORLD SCIENTIFIC PUBLISHING CO.; 2009. p. 205–11. doi:10.1142/9781848165632_0019.

41. Pfam: Family: Transferase (PF02458). https://pfam.xfam.org/family/PF02458. Accessed 22 Mar 2022.

42. Nelson DR. Cytochrome P450 diversity in the tree of life. Biochim Biophys Acta - Proteins Proteomics. 2018;1866:141–54. doi:10.1016/j.bbapap.2017.05.003.

43. Pfam: Family: UDPGT (PF00201). https://pfam.xfam.org/family/PF00201.21. Accessed 22 Mar 2022.

44. Phytozome. https://phytozome-next.jgi.doe.gov/. Accessed 22 Mar 2022.

45. TAIR - Home Page. https://www.arabidopsis.org/. Accessed 22 Mar 2022.

46. Consortium U. UniProt: the universal protein knowledgebase. Nucleic Acids Res. 2017;45: D158–69. doi:10.1093/nar/gkw1099.

47. UniProtKB. UniProtKB/Swissprot. https://www.uniprot.org/uniprot/?query=reviewed:yes. Accessed 22 Mar 2022.

48. Lallemand LA, Zubieta C, Lee SG, Wang Y, Acajjaoui S, Timmins J, et al. A Structural Basis for the Biosynthesis of the Major Chlorogenic Acids Found in Coffee. Plant Physiol. 2012;160:249–60. doi:10.1104/PP.112.202051.

49. RCSB PDB - 5YLW: CYP76AH1 from Salvia miltiorrhiza. https://www.rcsb.org/structure/5ylw. Accessed 29 Jun 2022.

50. Modolo L V., Li L, Pan H, Blount JW, Dixon RA, Wang X. Crystal Structures of Glycosyltransferase UGT78G1 Reveal the Molecular Basis for Glycosylation and Deglycosylation of (Iso)flavonoids. J Mol Biol. 2009;392:1292–302.

51. Katoh K, Standley DM. MAFFT Multiple Sequence Alignment Software Version 7: Improvements in Performance and Usability. Mol Biol Evol. 2013;30:772–80. doi:10.1093/molbev/mst010.

52. Larsson A. AliView: a fast and lightweight alignment viewer and editor for large datasets. Bioinformatics. 2014;30:3276–8. doi:10.1093/bioinformatics/btu531.

53. Yachdav G, Wilzbach S, Rauscher B, Sheridan R, Sillitoe I, Procter J, et al. MSAViewer: interactive JavaScript visualization of multiple sequence alignments. Bioinformatics. 2016;32:3501. doi:10.1093/BIOINFORMATICS/BTW474.

54. NCBI Multiple Sequence Alignment Viewer 1.22.0. https://www.ncbi.nlm.nih.gov/projects/msaviewer/. Accessed 2 Jul 2022.

55. Huang Y, Niu B, Gao Y, Fu L, Li W. CD-HIT Suite: A web server for clustering and comparing biological sequences. Bioinformatics. 2010;26:680–2. doi:10.1093/bioinformatics/btq003.

56. CD-HIT Suite. http://weizhong-lab.ucsd.edu/cdhit-web-server/cgi-bin/index.cgi?cmd=cd-hit. Accessed 29 Jun 2022.

57. Altschul SF, Gish W, Miller W, Myers EW, Lipman DJ. Basic local alignment search tool. J Mol Biol. 1990;215:403–10. doi:10.1016/S0022-2836(05)80360-2.

58. Altschul SF, Madden TL, Schäffer AA, Zhang J, Zhang Z, Miller W, et al. Gapped BLAST and PSI-BLAST: a new generation of protein database search programs. Nucleic Acids Res. 1997;25:3389–402. http://www.ncbi.nlm.nih.gov/pubmed/9254694. Accessed 22 Jun 2017.

59. Protein BLAST: search protein databases using a protein query. https://blast.ncbi.nlm.nih.gov/Blast.cgi?PROGRAM=blastp&PAGE_TYPE=BlastSearch&LINK_LOC=blasthome. Accessed 29 Jun 2022.

60. Pfam: Home page. https://pfam.xfam.org/. Accessed 29 Jun 2022.

61. T J, M P. Dotlet: diagonal plots in a web browser. Bioinformatics. 2000;16:178–9. doi:10.1093/BIOINFORMATICS/16.2.178.

62. Dotlet JS. https://dotlet.vital-it.ch/. Accessed 29 Jun 2022.

63. Huson DH, Scornavacca C. Dendroscope 3: An interactive tool for rooted phylogenetic trees and networks. Syst Biol. 2012;61:1061–7. doi:10.1093/sysbio/sys062.

64. Pearson WR, Lipman DJ. Improved tools for biological sequence comparison. Proc Natl Acad Sci U S A. 1988;85:2444–8. doi:10.1073/PNAS.85.8.2444.

65. LALIGN/PLALIGN local alignments. https://fasta.bioch.virginia.edu/fasta_www2/fasta_www.cgi?rm=lalign&pgm=pal. Accessed 29 Jun 2022.

66. Mauri M, Elli T, Caviglia G, Uboldi G, Azzi M. RAWGraphs: A visualisation platform to create open outputs. ACM Int Conf Proceeding Ser. 2017;Part F131371.

67. Plotly: The front end for ML and data science models. https://plotly.com/. Accessed 29 Jun 2022.

68. Lai C, Kunst L, Jetter R. Composition of alkyl esters in the cuticular wax on inflorescence stems of Arabidopsis thaliana cer mutants. Plant J. 2007;50:189–96.

69. Jumper J, Evans R, Pritzel A, Green T, Figurnov M, Ronneberger O, et al. Highly accurate protein structure prediction with AlphaFold. Nat 2021 5967873. 2021;596:583–9. doi:10.1038/s41586-021-03819-2.

70. Kriegshauser L, Knosp S, Grienenberger E, Tatsumi K, Gütle DD, Sørensen I, et al. Function of the HYDROXYCINNAMOYL-CoA:SHIKIMATE HYDROXYCINNAMOYL TRANSFERASE is evolutionarily conserved in embryophytes. Plant Cell. 2021;33:1472–91. doi:10.1093/PLCELL/KOAB044.

