## supplemental document 1 for "Seqrutinator: Non-Functional Homologue Sequence Scrutiny for the Generation of large Datatsets for Protein Superfamily Analysis"

### Supplemental Document 2: Technical Description Seqrutinator

#### Introduction.

Seqrutinator is a modular pipeline written in Python 2.7 to identify and remove protein superfamily sequences that likely either correspond to a non-functional homologue (NFH) or are incorrect, the latter due to either sequencing or gene modelling errors. Seqrutinator is made for large single-domain protein superfamilies that are nevertheless complex. As such it requires intermediate (>50 input sequences) or large datasets. This document describes in detail how the modules function, how the pipeline is made and which outputs are generated. Seqrutinator requires a single input file with the aligned sequence set in Fasta format and in addition trimmed at both the N- and the C-terminal end. The rationale for this as well as the biological fundamentals are described in the main manuscript. Seqrutinator was designed and tested on complete proteome-based datasets as well as on a control dataset that consisted of all SwissProt entries of the same superfamilies. It can most likely also be used for other datasets. In the main manuscript, we also describe how the origin and size of the dataset can affect its performance.

Seqrutinator is a flexible, modular pipeline. Here we describe how each module works as well as the required input and produced output. We describe how the pipeline is organized. Dependencies are described/listed in Materials and Methods, which includes shell commands that are used to run software packages such as MAFFT.

Seqrutinator starts with a check for sequences with non-IUPAC code and removes these. Each module is accessed by a number, according to the default pipeline 12345: 1: Short Sequence Remover (SSR) → 2 Non-Homologous Hit Remover (NHHR) → 3 Gap Instigator Remover (GIR) → 4 Continuous Gap Sequence Remover → 5 Pseudogene Remover (PR). Modules 2 to 5 need an MSA as input. To make it possible to construct customized pipelines, this is also required for module 1 even though it does not actually use the alignment. As such, Seqrutinator is best combined with another script, MUFASA, that mines and aligns protein superfamily sequences. In addition, each module ends with the construction of a novel MSA. Both MUFASA and Seqrutinator have been tested extensively using a Ryzen 7 16-thread CPU. All descriptions concern the default settings but many options to overrule default settings are described. As described in the main text of the manuscript, the user needs to trim the MSA that derives from MUFASA, typically using the sequence of a protein for which a structure has been resolved: The MSA should represent the mature protein and lack non-homologous terminal sequences. The sequence of that protein needs to be aligned by for instance MAFFT add.

Seqrutinator can be applied in batch using an additional script name SeqYNet.py.

#### <u>Multiple FASta Aligner</u>

MUFASA takes as active input a single HMMER profile (profileX.hmm, either custom made or from for instance Pfam). It then performs a *hmmsearch* of all fasta files that should be made available in separate subfolders that are located in the same folder as where MUFASA is executed. It collects all sequences from the fasta file that score above the sensitive HMMER inclusion threshold and it aligns them using

MAFFT G-INS-i. Both the unaligned (bug_profileX.fsa) and aligned sequences (bug_profileX.faa) are saved in the same directory as the original fasta file.

Command line: *python MUFASA.py -i <profileX.hmm>*
Additional options are described and can be acquired by *python MUFASA.py -h*

**Seqrutinator**

The Short Sequence Remover Module (1)

SSR opens the offered fasta file (*bug_profileX_3D.faa*) and determines the length *l* of the first sequence, which by default is the reference sequence. Next, it determines the length of each sequence and removes sequences < 0.65**l*, to bug_profileX_SSR_Removed.fsa, the remaining sequences are saved as bug_profileX_SSR.fsa, aligned by MAFFT G-INS-i and saved as bug_profileX_SSR.faa. The additional output consists of bug_profileX_SSR_removed.txt and contains the accession codes of the removed sequences.

Options for SSR include the change of the percentage for the length cut-off, pointing another sequence as reference or simply entering the length *l*.

Non-Homologous Hit Remover Module (2)

NHHR removes HMMER score outliers. It constructs a HMMER profile of the offered MSA bug_profileX_SSR.faa. It uses it to run a hmmsearch against the unaligned sequences and directly removes all sequences that score 0. It then uses the total scores of the sequences to perform a distribution analysis. In order to prevent removing short sequences (in case SSR does not precede NHHR in the applied pipe), sequences shorter than 65% of the reference sequence are excluded from this analysis. It determines the mean and the standard deviation.

Finally, it determines which sequences score below the threshold and removes these to bug_profileX_NHHR_Removed.fsa. The default cut-off is the three-sigma rule of thumb (also known as the 3σ rule: < mean minus three standard deviations). The remaining sequences are saved as bug_profileX_NHHR.fsa and aligned as bug_profileX_NHHR.faa. NHHR is not iterated since its objective is to remove sequences that are not homologous and will be removed in a single screening, under the assumption that they score low.

NHHR comes with optional threshold settings. Any value *alpha* can be used for the sigma rule. Alternatively, the cut-off can be based on interquartiles. The Inter Quartile Range or IQR is defined as the difference between the middle of the first half of the sequence scores and the middle of the second half of the sequence scores. This provides another measure that is often used to define outliers, such as the lower 1.5 * Inter Quartile Range (IQR) whisker (< Q1 – 1.5 x IQR). NHHR provides a histogram showing the score distribution with the cut-off that was applied as additional output.

Gap Instigator Remover (3)

GIR removes sequences that instigate large (>29) gaps as detected in the offered MSA bug_profileX_NHHR.faa. This comes with two uncertainties of which one is tackled automatically. Due to various reasons, a "sequence-specific insert" in general is never that specific that it instigates the same gap in all other sequences. Since the subsequence contains information that is used by MAFFT in the MSA reconstruction, residues and or subsequences from other sequences will be aligned to the sequence-specific insert. As such, we define the *majority gap column*, where majority means > 90% of the sequences. GIR determines which columns are majority gap columns and subsequently scans the MSA and determines the number of continuous regions of majority gap columns. It makes two counts. The *continuous gap score* resembles the local score: it sums but drops to zero when a normal column is encountered. The *combined gap score* resembles the global score: it sums and simply distracts each normal column that is encountered. The *continuous gap score* is used for the automated identification of GIR-NFH. GIR then removes the sequence that has the gap with the highest continuous gap score above the threshold to bug_profileX_GIRN_Removed.fsa. The remaining sequences are saved as bug_profileX_GIRN.fsa and aligned as bug_profileX_GIRN.faa, where N is the number of the iteration. Next, the procedure is iterated with bug_profileX_GIRN.faa as input. Hence, GIR is a one-by-one sequence remover with iteration.

GIR comes with optional settings for the cut-of: both the length of the continuous gap and the percentage of gaps for the definition of a *majority gap column* can be set. GIR provides plots for both the *continuous gap score* and the *combined gap score,* for each iteration. As is described in the main text, the *combined gap score* can be used for the manual removal of additional sequences.

<u>Continuous Gap Sequence Remover (4)</u>

CGSR removes sequences that have large continuous gaps as detected in the offered MSA bug_profileX_GIRN.faa. Since this gap can be the result of a family-specific deletion, gaps in *high gap columns* (>50%) are not considered. Given the high complexity of MSAs with sequences that have these large continuous gaps, these MSAs tend to be highly irregular. As such and as an extra precaution, CGSR determines all gap sizes and selects the gaps larger than 30 residues and in the upper 1.5 * IQR whisker (> Q3 + 1.5 IQR).

CGSR removes the detected sequences to bug_profileX_CGSRN_Removed.fsa. The remaining sequences are saved as bug_profileX_CGSRN.fsa and aligned as bug_profileX_CGSRN.faa, where N is the number of the iteration. Next, the procedure is iterated with bug_profileX_CGSRN.faa as input. Hence, CGSR is a controlled batch-sequence remover with iteration.

CGSR comes with optional settings for the determination of the cut-off, both the length of the continuous gap and the percentage of gaps for the definition of a *high gap column* can be set.

As a final remark, similar to what can happen to a sequence-specific insert, sequence-specific deletions can result in the misalignment of one or more residues that flank the deletion. As a result, such deletions can appear as split which impedes their detection. This problem is not tackled since the solution similar to that was used for GIR (combined gap score) tends to identify many loop regions and leads to many false positives.

<u>Pseudogene Remover (5)</u>

The pseudogene remover is the same outlier remover as NHHR but is iterated. It constructs a HMMER profile and runs hhmsearch against the unaligned sequences with the same default settings as NHHR. PR removes the sequences detected below the threshold to bug_profileX_PRN_Removed.fsa. The remaining sequences are saved as bug_profileX_PRN.fsa, aligned and saved as bug_profileX_PRN.faa, where N is the number of the iteration. This procedure is iterated with bug_profileX_PRN.faa as input. Hence, PR is a batch-sequence remover with iteration. It also tests if the final hmmsearch resulted in a normal score distribution.

PR comes with optional settings for the determination of the cut-off and provides a more elaborate graph than NHHR. It shows the actual score plot as well as a plot for the score-drops between two consecutive hits in the HMMER search (delta-score) and, additionally, four putative cut-offs. Besides the default $3\sigma$, it presents the $2\sigma$ threshold, the upper 1.5*IQR whisker (> Q3 + 1.5 IQR) and the major score-drop as putative cut-off threshold. Application of the major score drop corresponds to the idea that non-outliers (read functional homologues under functional constraint) evolve with similar evolutionary rates which results in a continuous HMMER score distribution whereas NFHs lack the constraint and will evolve faster and show much lower scores. In certain cases, the major score drop can detect this. Note however that the dataset supposedly contains sequences from various subfamilies that have somewhat different constraints and that this can also result in large score-drops. The more complex a superfamily is, the less reliable the idea of the largest score-drop is. Hence, the largest score drop should only be applied when it occurs at or near one of the other three thresholds.

<u>The pipeline</u>
Seqrutinator (seqrutinator_1.py) is a fully flexible modular pipeline. It consists of a main python script with all modules in a single script and some recurring algorithms in a python module (lib_seqrutinator_1.py). Minimal command is

*python seqrutinator_1.py -f <msa.faa>*

in which case it runs the default pipeline 12345 with the default cut-off thresholds as described. For any other mode of settings please apply the information below (python seqrutinator_v1.py -h):

usage: seqrutinator.py (-h) (-m M) (-f F) (-ali ALI) (-ref1 REF1) (-ref2 REF2)
(-BMGE BMGE) (-p1 P1) (-p2 P2) (-m2 M2) (-s2 S2)
(-a2 A2) (-m3 M3) (-p3 P3) (-aa3 AA3) (-p4 P4)
(-aa4 AA4) (-a5 A5)

optional arguments:
-h, --help show this help message and exit
-m M Pipelines
-f F Fasta file
-ali ALI MAFFT GINSI (0), MAFFT Global (1), MAFFT Auto (2), FAMSA (3)

-ref1 REF1 Name of input reference sequence
-ref2 REF2 Users input reference sequence length
-BMGE BMGE BMGE deactivated (0), BMGE > 0 is h option for BMGE
-p1 P1 Percentage of sequence length coverage for SSR
-p2 P2 Percentage of sequence length coverage for NHHR
-m2 M2 HMMER Score (1) or TCS (2) method for NHHR
-s2 S2 Mean - alphaSD (1) or Q1 - 1.5IQR (2)
-a2 A2 Alpha for NHHR
-m3 M3 Method one-by-one (1) or batch (2) for GIR
-p3 P3 Percentage of gaps to define *majority gap column* for GIR (>= VALUE)
-aa3 AA3 length of aa window of continuous gap columns for GIR
-p4 P4 Percentage of gaps to define *high gap column* for CGSR (>= VALUE)
-aa4 AA4 length of aa window of continuous gap columns for CGSR
-a5 A5 Alpha for mean - alphaSD (3 is recommended as default option and

2.35 for normal distributions)

Example:
*python seqrutinator_v1.py -f sample.fsa -m 1324 -p1 85 -m2 2 s2 2 -a2 5 -m3 2 -p3 80 -aa3 35 -p4 85*
Runs seqrutinator in the order SSR-GIR-NHHR-CGSR, hence it will remove sequences <85% of the length of the first sequence in *sample.fsa,* it will then iteratively remove sequences that instigate gap regions of 35 (where a gap column has more than 80% gaps) in batch mode, next it removes sequences with gap regions of > 30 where 85% of the sequences are allowed to also have a gap at the same column. Do note that the latter is actually not a good idea since it can result in the exclusion of a sequence that has a large gap in the MSA where many sequences actually have gaps.

At the end of each module seqrutinator performs a BMGE test that determines the number of columns before and after trimming and saves the numbers in a tsv file. The rationale of this is explained in the main text of the manuscript.
