## supplemental document 2 for "Seqrutinator: Non-Functional Homologue Sequence Scrutiny for the Generation of large Datatsets for Protein Superfamily Analysis"

### Supplemental Document 2: Sequence Recovery Analysis Seqrutinator.

The recovery analysis of the sequence sets obtained from SwissProt, as described in the main text, indicates that automated sequence scrutiny comes with only a few false positives and that it is sensitive since it detected GT28 sequences, which form a superfamily related to the UGT superfamily. Here we analyse the output of the 16 proteome datasets in order to strengthen and expand the recovery analysis. Our approach consisted of identifying the conservation of traits among removed sequences. The main underlying thought is that subfamilies can be represented differently in complete proteomes than in a SwissProt sequence set. In the absence of orthologues, specific paralogues will stand out and may be removed from complete proteomes. As such, for each superfamily, we combined the sequences removed by either GIR, CGSR or NHHR and PR (combined) from the 16 complete proteome scrutinies to make three sets of sequences per superfamily. This allows for cluster analysis in order to detect conserved inserts, conserved gaps or distant subfamilies, respectively. In addition, we analysed the output of the more strict NHHR module in order to see if a more strict threshold would lead to the identification of similar homologues (i.e. the paralogues that are removed from the complete proteome datasets). We projected that sequences detected as outliers in the complete proteome scrutiny may be detected in a SwissProt sequence set only when a more stringent cut-off threshold is applied .Another reason for applying a more strict NHHR cut-off lies in the fact that NHHR did not detect false positives from the SwissProt sequence sets and that a more strict cut-off threshold may improve the results in terms of sensitivity. These analyses are described here in order to show Seqrutinator performance as well as how recovery analyses are most easily performed.

#### The outlier cut-off threshold is sufficiently strict

We analysed the output of pipeline pipe 12a345, which applies the more strict 2σ rule to NHHR. In the BAHD case, the strict NHHR module detected three curated SP entries. These entries, Uniprot identifiers Q39048, Q9LIS1 and Q9SVM9, correspond to entries belonging to the eceriferum or CER subfamily, a distant BAHD subfamily characterized by the absence of the DFGWG motif and potential ER localization. Their molecular function is unknown but they have been implicated in the mechanism of cell wall waxes elongation (68).

A total of 35 putative BAHD encoding sequences (NHHR 5, PR 30) were removed from the 16 proteomes by the default pipeline. We performed a CD Hit clustering and all 14 sequences from CD Hit cluster 0 as well as all three sequences from CD Hit cluster 2 were, by means of phylogeny and HMMERCTTER clustering, shown to be part of the CER subfamily (See Figure SD2-1). This confirms the hypothesis that the different representations of certain subfamilies in complete proteomes may result in a different Seqrutinator performance. Note that not all CER subfamily members were removed but that the removed and reintroduced sequences appear at relatively large evolutionary distances, which explains they were detected as outliers. The result indicates that the 3σ threshold is sufficiently strict, contrary to what the results on the SwissProt dataset suggested. At least for this case, the default setting results in a balance between sensitivity and precision. The fact that the CER subfamily lacks the DFGWG motif suggests that these BAHD homologues are in the twilight zone of the superfamily. The removal of the other sequences was correct since they hit no or other Pfam profiles.

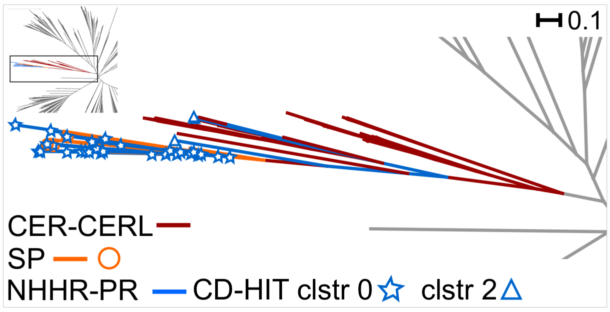


***Figure SD2-1: NHHR false positives cluster with the distant CER subfamily****. Phylogeny of SwissProt-accepted and 17 BAHD sequences removed by NHHR or PR (CD Hit clusters 0 and 2). Reintroduced sequences fall in a clade that forms a 100% P&R cluster that includes a subclade with all SwissProt CER entries. No other annotated sequences were found in the clade.*

A similar analysis performed for CYP shows similar results. This analysis is described in more detail in order to demonstrate the additional output of Seqrutinator. The output of Seqrutinator includes a log that indicates the major events as well as a number of files with graphs that can assist in the analysis of the sequence scrutiny (See examples in figure SD2-2, A to C default pipeline, D to F 12a345 pipeline). The output also includes an archive with the results of the trimming analysis that shows if sequence scrutiny has resulted in an improved MSA, as is discussed in the main text. The plots provide visual information on the performance of the modules. The first plot shows the HMMER score distribution of the NHHR analysis. Figure SD2-2A shows that with the default setting no sequences scored below the cut-off of 3σ (Mean-3SD), whereas Figure SD2-2D shows 17 sequences were removed by NHHR with the more stringent 2σ (Mean-2SD) setting. Figures SD2-2C and 2F show HMMER score plots provided by PR at the end of these scrutinies. These show the alternative cut-offs of 2σ and the interquartile HIQR (Q3+1.5IQR).

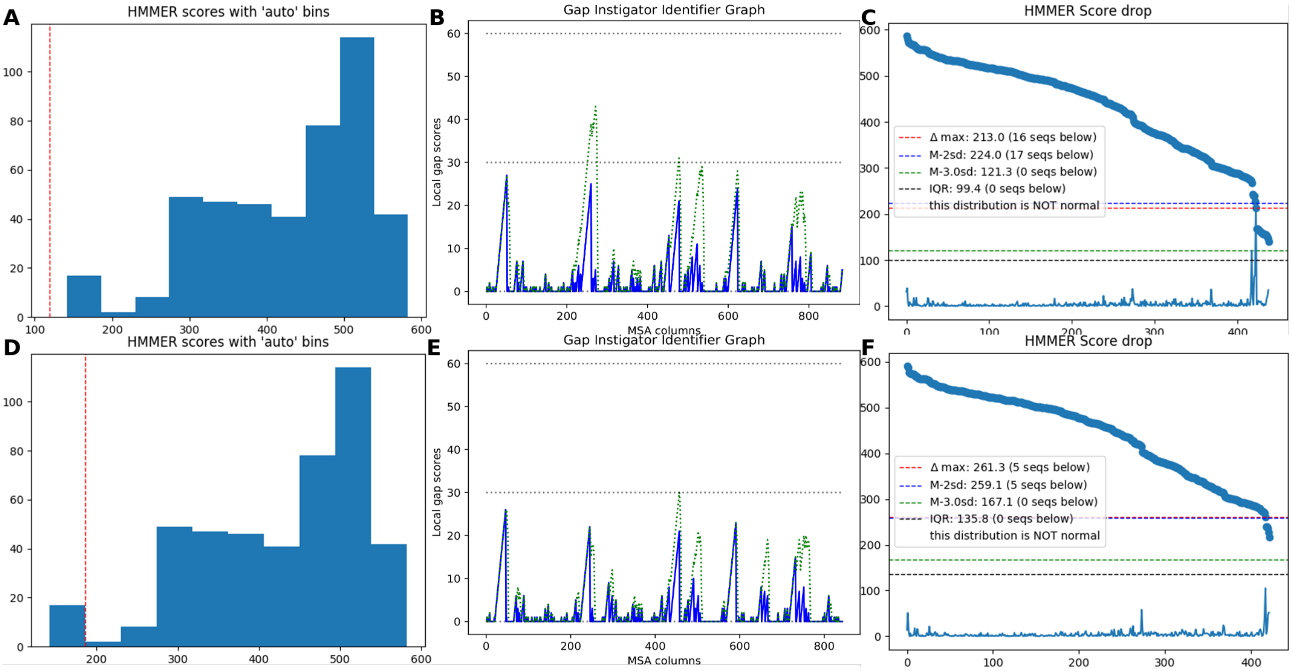


***Figure SD2-2: Recovery SwissProt CYP case. A and D:*** *Score distribution histograms of NHHR modules on CYP, Curated SwissProt Sequence set, with cut-off (red dashed line) at 3σ and 2σ, respectively.* ***B and E:*** *Gap score plots of GIR modules applied to sequence sets following NHHR with 3σ and 2σ, respectively. The blue line shows the applied continuous gap score and the dashed green line the more strict combined gap score (see also main text).* ***C and F:*** *Score plots of PR modules applied to sequence sets following CGSR, after NHHR with 3σ and 2σ, respectively. Indicated are putative cut-off thresholds. The blue line graph is the plot that shows the score drop between respective hit scores.*

In addition, they show the largest score-drop (Δmax), which sometimes appears as a helpful guide when it concerns outliers. Interestingly, Figure SD2-C shows outliers in two groups with similar scores. The lower cluster has 16 of the 17 sequences that NHHR removes with the 2σ setting and encode allene synthases. The 17$^{th}$ sequence removed by NHHR 2σ is a partial sequence and is normally removed by CGSR. The second group of five sequences corresponds to C-22 sterol desaturase. These are all functional homologues and are by default not removed, confirming again 3σ as the default threshold. However, this example does show how Δmax can assist. Application of the 2σ cut-off would not only remove the 16 allene synthase homologues but also one of the five 22 sterol desaturases. Applying the Δmax would remove the allene synthase sequences only.

Figure SD2-2B and Figure SD2-2D are reports from the GIR module as applied again in the default and strict NHHR pipeline. They show the lengths of instigated, single continuous gap regions that are used to remove eventual NFHs. The graph also includes a more sensitive "combined gap" score that corresponds conceptually to the identification of large putative gaps that may have become split by one or a few incorrectly placed residues. The log indicates the existence of such putative combined gaps and the user can choose to remove these upon visual inspection. Two such combined gaps were identified in the default pipeline. Interestingly, these large combined gaps disappeared when we applied the stringent setting in the NHHR module which indicates that one or more of the 17 sequences that were removed by NHHR at 2σ instigate a large gap. The fact that the same sequences are identified as putative rather than real NFH shows the pipeline's versatility and robustness.

Seqrutinator identified 76 outliers in the 16 proteomes, either from NHHR (15) or PR (61) which were combined for the recovery analysis. First, batch Pfam analysis showed that 17 outliers are not CYP since they hit other or no domains. The remainder of the sequences was then analysed by BLAST and this did identify 49 false positives that all derive from the PR module. Of these, 48 are part of the allene oxide synthase subfamily and one is a C-22 sterol desaturase. Hence, again, the results of a strict NHHR scrutiny of SwissProt appear to predict problems with the separate proteomes. Also these results confirm that the 2σ is too strict since 3σ removed a number of distant CYP sequences.

The remaining ten outlier sequences were correctly removed. One sequence concerned B3LF83.1 is a partial sequence also present in SwissProt (See Table 1 of main text), whereas five sequences showed either or both very low identity or cover to other SwissProt sequences, which suggests these concern pseudogenes. A single case contains a partial repeat and also most likely corresponds with a pseudogene. We also found two sequences yielding truncated alignments, which suggests these sequences derive from incorrect gene models. The last relevant positively detected NFH shows the most similarity to a Monocopper oxidase-like protein SKS2 which is not a CYP, despite the fact that Pfam accepts it as CYP with an E-value of 6E-22.

An alternative and faster analysis of the CYP outliers is by CD Hit cluster analysis. The first cluster contained 50 sequences among which the 48 sequences that were annotated as allene oxide synthase (or related). The 49$^{th}$ sequence is a partial allene oxide synthase and the last is the sequence of an allene oxidase pseudogene with a partial repeat in the C-terminus. BLAST analysis of this sequence cannot align it properly but it aligns quite well in the MSA of the subset of sequences. The likely functional C-22 sterol desaturase would have been missed in this analysis.

Perhaps the most astounding part of the recovery analysis is that of the Glycosyl Transferase 28 (GT28) enzymes involved in the biosynthesis of steryl and acyl steryl glycosides, of which two sequences were identified from the SwissProt dataset. A total of 61 GT28 sequences were identified from the separate proteome sets, interestingly, not only by NHHR (6) and PR (10) but also by GIR (12) and CGSR (33). These sequences were aligned to the final MSA of accepted sequences. All GT28 sequences were substantially different from the UGT sequences but also showed mutual similarity. The MSA (excerpt showing large inserts in Figure SD2-3A) was used to reconstruct a phylogenetic tree. Figures SD2-3B and 3C show that the removed GT28 sequences cluster together in a clade at a large distance. Interestingly, the clade appears as a subclade from a clade with four sequences, one from *Amborella trichopoda* and three from *Physcomitrela patens.* Two are annotated as steryl-glycosidases and all four sequences are not identified in a Pfam scan using the gathering threshold. When we applied a sensitive cut-off of E-value 1, Pfam detects them as GT28. Figure SD2-3A is an excerpt that shows how GT28 sequences but not the other four sequences from the mentioned clade disturb, among others, the PSPG motif. Besides that the sequences instigate large gaps, they have large gaps, which explain why some of them were removed by GIR or CGSR, rather than NHHR.

The four sequences represent an example of twilight zone sequences.They appear as UGT in the tree and they have a perfect PSPG motif but they are also quite well related to GT28. We checked if other sequences, similar to the four twilight zone GT28-UGTs, had been removed from the non-seed plant sequence set. We identified four of the 61 removed GT28 sequences. Again, the default setting appears as correct. This also shows why there is no benchmark dataset for sequence scrutiny. Besides that it is hard to show a sequence is not functional, there are often no clear criteria for sequence classification.

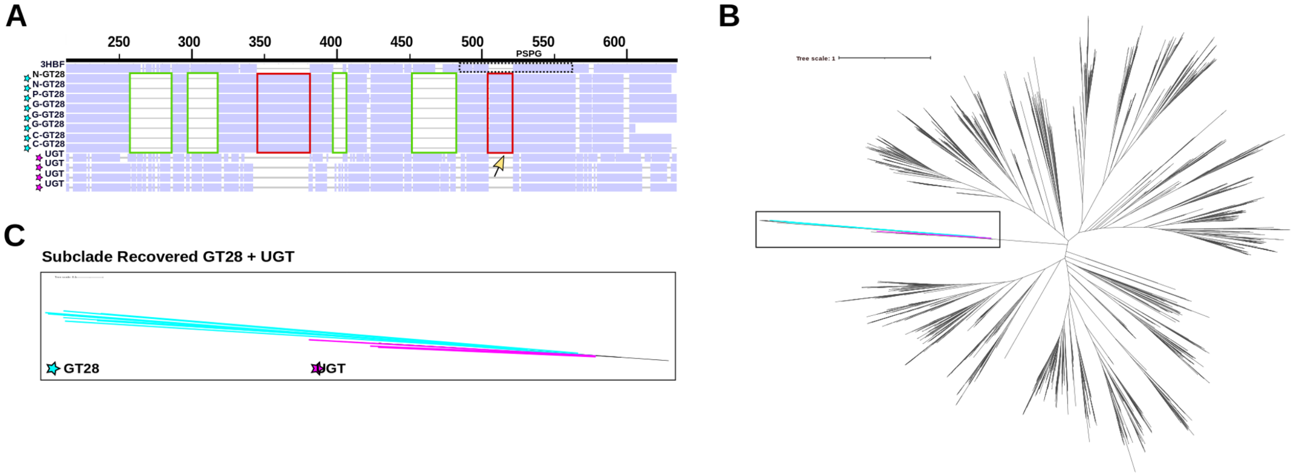


***Figure SD2-3. Removed Glycosyl Transferase 28 sequences are not UGT and are real positives. A:*** *Excerpt of an MSA with GT28 sequences removed by the NHHR (N), PR (P), GIR (G) or CGSR (C) module and aligned to the final UGT MSA. The GT28 sequences have inserts (red boxes) among which a 17 residue insert at position 505 (yellow arrow) that disturbs the PSPG motif (black dotted box). GT28 sequences also instigate large gap regions (green boxes). Four sequences demonstrated below the GT28 sequences fall in the same clade (see panels B and C). 3HBF (R) is the reference sequence for UGT.* ***B:*** *Phylogeny of accepted UGT and reintroduced GT28 sequences.* ***C:*** *Detail of GT28-UGT clade with GT28 sequences in light blue and four UGTs in magenta.*

We also analysed the other sequences identified by NHHR or PR. All seven sequences clustered in cluster 1 are epimerases (PF01370). Cluster 3 contained a mere three sequences, two from potato and one from tomato. Hence, we envisaged these might represent a taxon-specific subfamily. Indeed highly similar sequences from other Solanum species as well as Solanaceae species as *Capsicum annuum* (bell pepper) were identified performing a BLAST against Reference Proteins. A sudden drop of identity from ~80 to ~58% occurred from a hit against *Nicotiana tabacum* (tobacco, another Solanaceae) to the next hit from *Ipomea nil* (morning glory, a Convolvulaceae from the order of Solanales). This suggests these sequences, which do not appear to have any other problems when aligned against the SwissProt MSA, represent a taxon-specific subfamily. As such, this shows that screening complete proteome-derived sequence sets may result in false positives, which can be analysed quickly given the output of Seqrutinator.

All together these analyses show that Seqrutinator modules NHHR and PR remove real outliers but also that on occasion they remove sequences that encode functional homologues belonging to distant subfamilies. The default cut-off of 2$\sigma$ appears to result in a good balance in terms of precision and recall, particularly when a recovery analysis is included.

**The GIR module Identifies False Positives in the BAHD case but not in the CYP and UGT cases.**

A total of 54 candidate BAHD sequences were removed by GIR in the default pipeline. All sequences presented significant hits to the (BAHD) Transferase domain, as determined by hmmscan. Three sequences also presented partial but significant hits to other families (TIR PF01582.23, Flavodoxin PF00258.28 and Peptidase_M14 PF00246.27), which explains why these instigate a large gap in an otherwise BAHD alignment. Many of the removed sequences presented more than one significant hit with the (BAHD) Transferase domain, hence they may present internal duplications. We analysed these sequences using LALIGN local alignment.

We found three scenarios corresponding to most of the sequences removed by GIR. Figure SD2-4 depicts examples for each scenario. In scenario 1, the (BAHD) Transferase model appears as broken, but sequences actually show at least one large single repeat. As such the sequences hit part of the model twice leading to total scores above the outlier threshold of the earlier NHHR module. Local alignment with LALIGN finds internal duplications with E-value < 0.0001, suggesting recent duplication events or errors made during sequencing, assembling or gene modelling. In scenario 2, part of the (BAHD) Transferase profile cannot detect its counterpart in the query sequence. This is combined with a repeat with a rather high E-value. This repeat is repeated, as is shown by the parallel secondary alignments, by which the subsequence is also relatively long, which explains also these sequences instigate a gap. In scenario 3, the model is typically well covered and no internal duplications are found (not shown).

The sequences were also subjected to CD Hit clustering. The largest cluster has 14 sequences depicting broad taxonomic distribution. The other clusters showed rather strict taxon distributions. The second cluster has eight sequences from *P. taeda* and *O. sativa*, the third is a cluster with six sequences from *P. patens*, *M. polymorpha* and *P. taeda* and the remaining clusters with four, three and twice two sequences all derive from *P. taeda* sequences. Since the dataset affects the performance, we then decided to include the 14 sequences from the largest cluster to the accepted SwissProt sequence set and apply the GIR module.

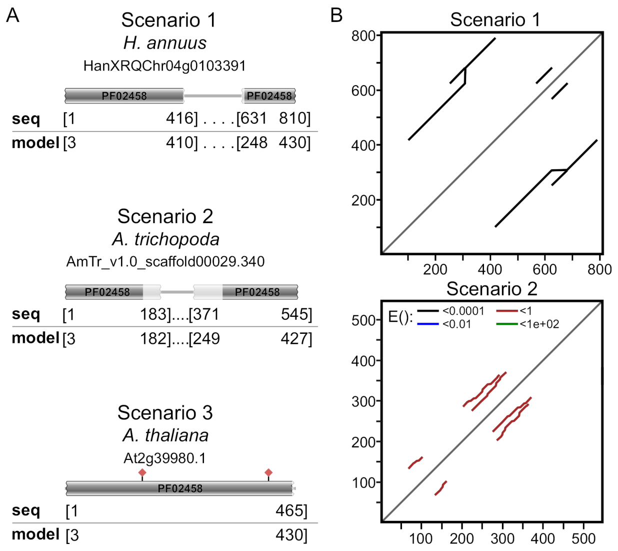


***Figure sd2-4: Different types of repetitions instigate gaps. A:*** *Sequences removed by GIR were analysed with hmmscan and mostly correspond to one of three scenarios. Shown are paradigmatic sequences (ids indicated) for each scenario. "seq" and "model" indicate the start and end positions of the hits of the query and the (BAHD) Transferase model as determined by Pfam. Red squares indicate conserved motifs.* ***B*** *Dotplot representation of LALIGN self-alignment. Secondary alignments indicate sequence repeats (E-value according to colours).*

Now only three of the sequences, which present either scenario 1 or 2, were removed. An inspection of the output MSA reveals that the 11 retained sequences have an insert on a conserved position (~420-480, Figure SD2-5A). One of the sequences was used for structure modelling using Alphafold (69) and the obtained model showed that the position is surface exposed.

Next, we reconstructed a phylogeny including the 11 sequences, and performed HMMERCTTER clustering. Ten of the recovered sequences fall in the same subgroup, part of a cluster that is 100% P&R (Figure SD2-5C). These sequences correspond to scenario 3. The 11$^{th}$ sequence (from *P. taeda*) appears as a sibling orphan of this clade. We conclude these 11 sequences form a second example of a possible incorrectly removed set of sequences.

The third CD Hit cluster has five entries from gymnosperm *P. taeda* and two entries from bryophytes *P. patens* (Pp3c2_29140V3.1.p) and *M. polymorpha* (Mapoly0003s0277.1). Both bryophyte and two of the *P. taeda* homologues have a large insert in the central region of the protein when aligned to the SwissProt dataset using MAFFT add (Figure SD2-5B). However, given that some of the inserts are short, it is not clear if the inserts are homologous. Four of the five *P. taeda* sequences have a large N-terminal extension which strengthens the idea that the inserts of the two *P. taeda* homologues are not monophyletic with the inserts of the gymnosperm sequences. We applied phylogeny in order to show how these sequences evolved. *P. patens* and *M. polymorpha* fall in the cluster with enzymes with hydroxycinnamoyl transferase (HCT) activity where the entries from *P. taeda*, whether with or without the insert, belong to a distant subfamily (Figure SD2-5D). Hence, although in many cases simple alignments can shed light on certain events, we must take care not to fall into the trap of parsimony. Both the CD Hit clustering and the perfect alignment of the inserts of two of the *P. taeda* and the two bryophyte entries suggest homology of the insert. Phylogeny clarifies however that most likely it concerns a hot spot of acceptable inserts. Both bryophyte homologues have been demonstrated to have HCT activity (70). Interestingly, *in vitro* HCT activity and even substrate preference is maintained in a mutant enzyme of the *P. patens* homologue in which the region corresponding to the insert has been deleted. This further corroborates the hypothesis of the hot spot for acceptable inserts. This is another example of where the assumption that large inserts are fatal for functionality appears incorrect.

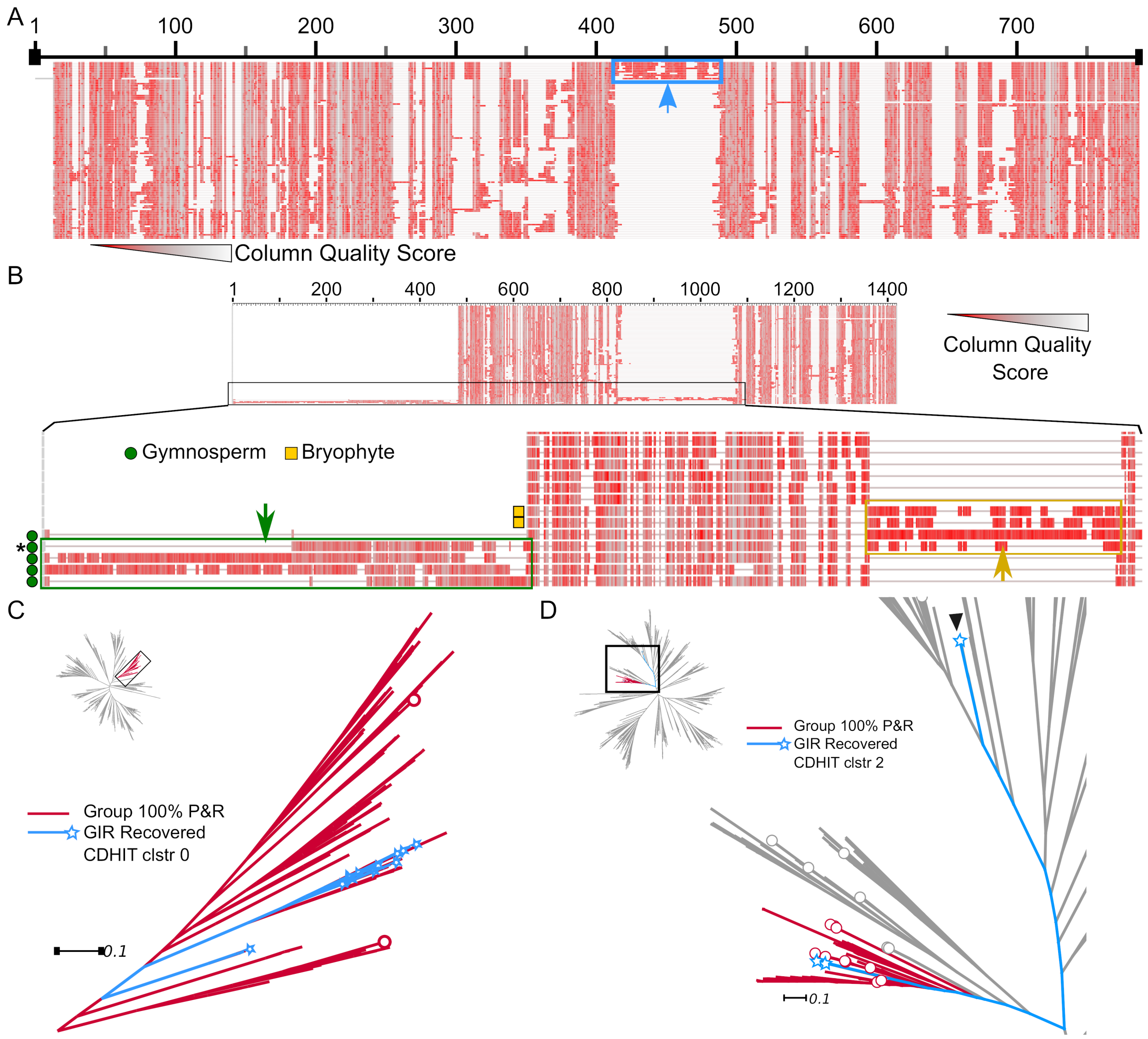


***Figure SD2-5: Recovery analysis of BAHD sequences removed by GIR. A:*** *MSA excerpt showing 11 sequences removed by GIR (top) instigate a large gap in the MSA of accepted SwissProt entries (~420-480, delimited by the box and indicated by an arrow). Column quality score is depicted according to the shown scale.* ***B:*** *MSA excerpt showing seven sequences removed by GIR and grouped in the third CD Hit cluster aligned to the SwissProt entries MSA. Sequences from* P. taeda *and bryophytes are indicated with green circles and yellow boxes, respectively. Four sequences, two from bryophytes and two from* P. taeda *instigate a large gap (yellow box) whereas four of the five* P tadea *sequences have a large N-terminal extension (green box). The star indicates a sequence with both an N-terminal extension and insert.* ***C:*** *Phylogeny of accepted SwissProt BAHD sequences and the 11 false GIR positives from CD Hit cluster 0 (shown in blue and with a star).* ***D:*** *Phylogeny of accepted SwissProt BAHD sequences and three false GIR positives from CD Hit cluster 2 (shown in blue and with a star). The two sequences from bryophytes* P. patens *and* M. polymorpha *fall in the 100% P&R clade with HCTs whereas the single* P. taeda *with only the insert (also indicated with an arrowhead) falls in a different clade.*

CD Hit clustering of the 112 putative CYP sequences that were removed by GIR yielded few clusters, the largest containing a mere 10 sequences. Nine of these sequences instigated different gaps of 30 or more residues in the MSA made of that cluster (Figure SD2-5A). Thus, although the sequences cluster, they differ in the gap they instigate. The tenth sequence instigated a gap of only 18 residues. However, when aligned this sequence to the SwissProt alignment (following Seqrutination) a gap of exactly 30 residues was instigated (Figure SD2-5B). Hence, on the one hand, also this sequence is unlikely to encode a functional homologue

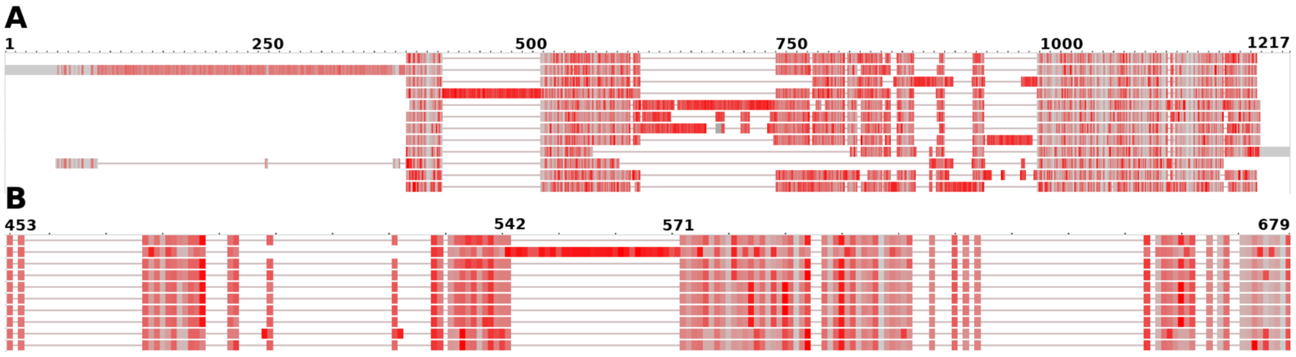


***Figure SD2-6: Recovery analysis of CYP sequences removed by GIR. A:*** *An MSA of the largest cluster of sequences obtained from GIR module of CYP case. Nine of ten sequences have inserts that result in a large gap in most of the other sequences.* ***B:*** *The tenth sequence shows a gap of exactly 30 residues when aligned to the MSA of the accepted SwissProt sequences as shown in B excerpt of MSA with 440 sequences 910 columns. The reference sequence 5YLW is shown on top of both A and B.*

and was correctly removed. On the other hand, it does point out that although Seqrutinator is objective it is sensitive to the data that are analysed. The other CD Hit clusters contained only a single pair of sequences with a homologous insert that appeared to be an internal repeat. Hence, in the CYP case, GIR did not yield a single false positive.

In the case of the UGTome, the largest cluster contained seven GT28 sequences, as described above. The second and third cluster contained six and five sequences respectively. Alignment consistently showed that the gaps that were instigated are not conserved and, interestingly, contain only sequences from *P. taeda*. Hence, we conclude that all sequences removed by GIR are most likely real non-functional homologues.

**No clear false positives could be recovered from the CGSR datasets**

Most of the NFH sequences are partial and most were removed by SSR, even using the safe setting of 0.65 times the length of the user selected reference sequence. In most cases, also larger sequences will likely not be functional. In many cases a more stringent setting can be applied upon criteria that are to be set by the user. In this study we chose a safe setting such that a recovery screen was not required for SSR removed sequences. Hence in the CGSR recovery screen we expect to identify mostly sequences that are incomplete at the N- and/or the C-terminal.

CD Hit Clustering of the 740 putative CYP sequences that were removed by CGSR yielded seven clusters of more than 20 sequences. The sequences of these clusters were aligned in the presence of the general reference sequence for CYP (PDB identifier 5YLW). Alignment by MAFFT tends to result in ordered MSAs in which often subfamilies can be visually identified. However, not a single subfamily could be detected. We then ordered the sequences such that:

1 Sequences that appear to lack N-terminal subsequence are at the top, ordered by the length of the missing subsequence. Note that these may also lack C-terminal subsequence.

2 Sequences that do not lack N-terminal but do lack C-terminal subsequence are at the bottom, ordered by the length of the missing subsequence.

3 The remainder of the sequences were ordered by the position of the first large (>30 aa) gap and occur as block in the middle.

Not a single subfamily with conserved gap was identified. Figure SD2-7 shows as an example the aligned sequences of the largest CD Hit cluster. As expected there are many sequences that lack part of either or both the N- or the C-terminus. Approximately 50 sequences have both the N- and the C term intact but have an internal gap. None of the internal gaps appears however as conserved. Interestingly a number of sequences also show intermediately long inserts.

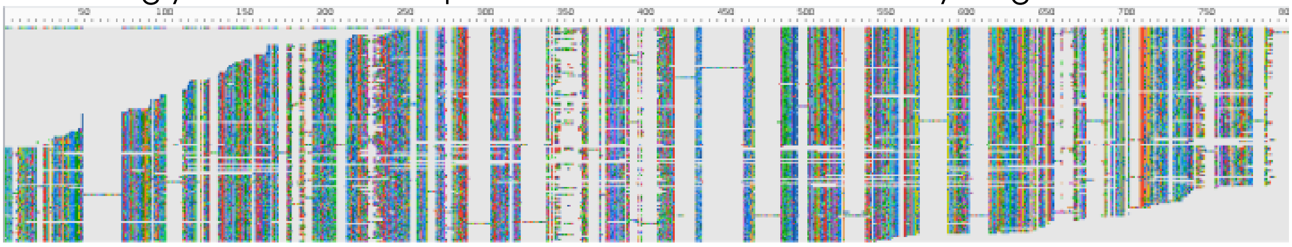

***Figure SD2-7: MSA of largest cluster of CGSR removed sequences.*** *A total of 133 sequences that was rejected by CGSR and clustered by CD HIt was aligned with general reference 5YLW (top). Next they were ordered based on lacking N-terminal or C-terminal sequences. Approximately 50 sequences with both the N- and C-termini intact were ordered by the position of the first large gap.*

A similar analysis was performed with the UGT datasets. We analysed the eight largest CD Hit clusters. Most sequences lack N-terminal sequence, many lack C terminal sequence and the remainder of the sequences all has large gaps which position was not conserved.
