## supplemental figure 2 for "Seqrutinator: Non-Functional Homologue Sequence Scrutiny for the Generation of large Datatsets for Protein Superfamily Analysis"

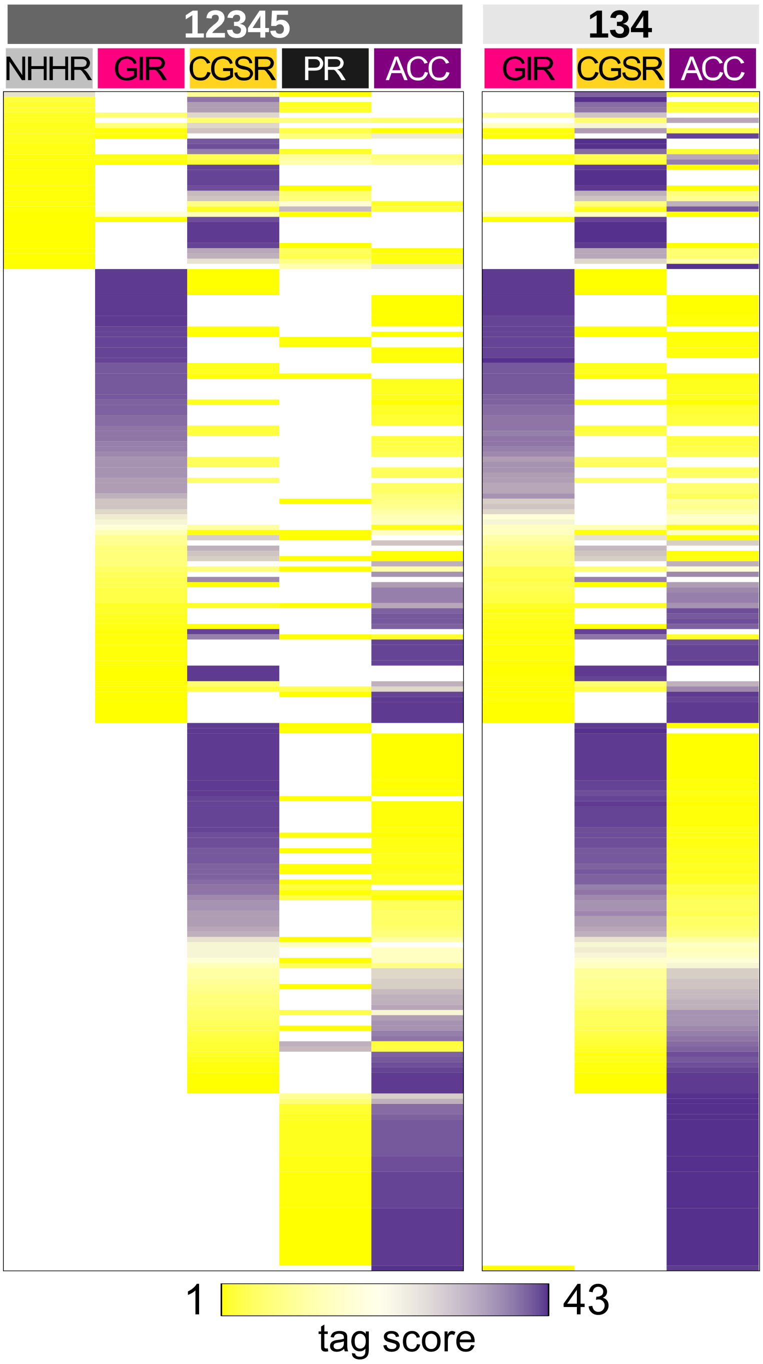


***Supplemental Figure 2: Sequences can be removed by different modules depending on the dataset.*** *Tag scores of 227 twilight-zone sequences for NHHR, GIR, CGSR or PR modules and final acceptance for pipelines 12345 (right) and 134 (left). Tag score according to heat map, white spaces indicate tag score of 0.*
