## supplemental figure 1 for "Seqrutinator: Non-Functional Homologue Sequence Scrutiny for the Generation of large Datatsets for Protein Superfamily Analysis"

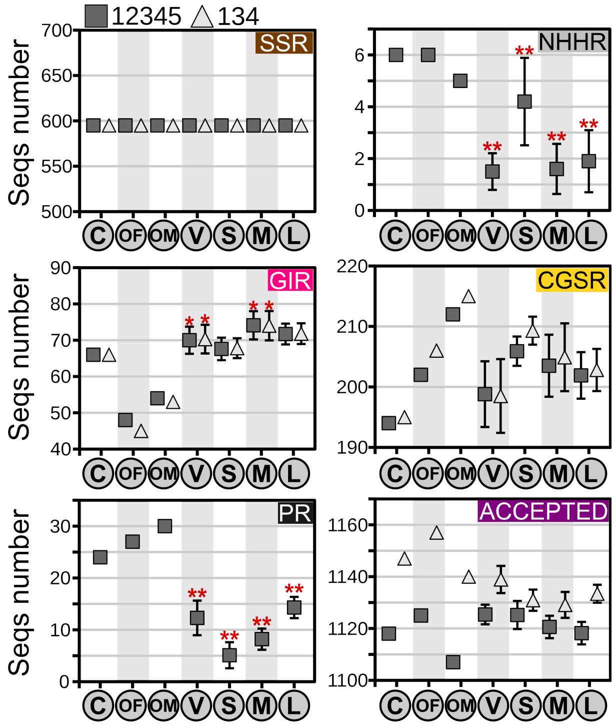


***Supplemental Figure 1: Netto Effect of Dataset on BAHD Sequence Removal.*** *The number of sequences removed by each module in either the default pipeline (12345) or the pipeline without outlier removers (134) was determined. C: Combined dataset (All sequences); OM: MAFFT-G-INS-i aligned original sequence sets; OF: FAMSA aligned original sequence sets; V Variable; S: Small; M: Medium; L: Large (L) sizes. For V, S, M and L ten randomized datasets were tested, average and s.d. are shown. Coefficient of variation, CV is indicated when >10% (**) or >5% (*).*
